# Structural Insights into the Expanded Substrate Scope of a Cysteine-Dependent Hydrolase from *Rhodococcus opacus* IITR-PK for Polyethylene, Polypropylene, and Polystyrene Biodegradation

**DOI:** 10.64898/2026.10.07.757487

**Authors:** Shubham Aggarwal, Saurabh Singh, Arpan Kayastha, Divya Aggarwal, Kuldeep Jangid, Mohammad Shakib, Deepika Singh, Ashwani Kumar Sharma, Pravindra Kumar

## Abstract

Microplastic pollution from homopolymers such as polypropylene (PP), polystyrene (PS), and polyethylene (PE) poses growing health and environmental concerns, highlighting the need for robust polymer-degrading biocatalysts. We previously identified a naturally occurring cysteine-dependent hydrolase (CsdH) from *Rhodococcus opacus* IITR-PK, featuring a unique Cys160–Lys127–Asp39 catalytic triad and providing a novel mechanism for polyethylene terephthalate (PET) degradation. In this study, we have determined crystal structures of CsdH in complex with PP, PS, PE, and styrene and demonstrated the potential of the enzyme to directly degrade these homopolymers into their respective low molecular weight degradation products using LC-MS. Structural analysis reveals that the enzyme retains a conserved principal access tunnel while undergoing subtle substrate-dependent changes to accommodate different substrates. The enzyme exhibited broad plastic degradation activity with the highest weight loss observed for PE, followed by PP and PS. SEM analysis reveals surface roughness, cracks and grooves, confirming polymer degradation. Further, these results corroborate through FTIR and TGA analysis. These findings establish a strong foundation for enzyme engineering and aim at developing efficient microplastic degradation.

## 1. Introduction

Synthetic plastics have revolutionised modern life due to their durability, low weight, chemical resistance and low manufacturing cost, which has resulted in their widespread production and consumption (Macheca et al., 2024). The world produces approximately 400 million metric tonnes of plastic annually, and this figure is anticipated to increase over the next few decades (J. Qiu et al., 2024; Srivastava et al., 2025; Verma et al., 2026). Three types of plastic, polyethylene (PE), polypropylene (PP) and polystyrene (PS), constitute around 60 % of all plastic produced (Kunwar et al., 2016; Meran et al., 2008). The polymers are used extensively in packaging, building, automotive, medical gadgets, textiles and consumer goods (Thadepalli, 2022). Overall, their excellent physicochemical stability has led to the continual buildup of plastic debris in terrestrial and aquatic environments, where they are persistent for decades and are progressively fragmented into microplastics and nanoplastics (Hossain and Engelhardt, 2025). These persistent pollutants have now been discovered in soils, freshwater systems, marine ecosystems, air aerosols, food chains and even human tissues, leading to major environmental and public health implications (Kayastha et al., 2026). Polyolefins such as PE and PP possess highly stable C–C backbones and lack hydrolysable functional groups, in contrast to biodegradable polyesters, which make them highly resistant to microbial breakdown (Pérez-García et al., 2025). The aromatic backbone of PS also leads to enhanced chemical stability and hydrophobicity, which in turn decreases microbial colonisation and enzymatic assault (Ho et al., 2018). Hence, the traditional waste management, such as landfilling, incineration and mechanical recycling, are still the dominating disposal methods (Hamer, 2003). However, these systems suffer from major restrictions in greenhouse gas emissions, creation of secondary pollutants, loss of polymer quality after repeated recycling, and low economic viability. Hence, eco-friendly biological technologies to turn plastic waste into reusable chemical feedstocks have gained great attention as promising options when considering the circular plastic economy (Nikolaivits et al., 2021). Microbial plastics degradation is largely mediated by extracellular or cell-associated enzymes that perform polymer oxidation or depolymerisation, which generate smaller oligomers and monomers that can then enter into core metabolic pathways (Singh et al., 2023; Srivastava et al., 2025). Significant advances have been made in the knowledge of the enzyme-based polyethylene terephthalate (PET) degradation with the discovery of PETase and MHETase from *Ideonella sakaiensis* (Maity et al., 2021). Protein engineering has been widely applied to enhance the catalytic efficiency and thermostability of PET-degrading enzymes, aided by structural studies (Son et al., 2019). PET biodegradation is one of the most developed fields of enzymatic plastic recycling at present (Singh et al., 2024a). In contrast, the enzymatic degradation of carbon-carbon backbone polymers is poorly understood since there are no hydrolysable ester bonds and the activation energy for cleavage of carbon-carbon links is quite high (Bher et al., 2022). Therefore, only a limited number of bacterial strains, microbial consortia, oxidoreductases (Danso et al., 2019), laccases, alkane hydroxylases (Liu et al., 2026), peroxidases (Alharbi et al., 2026) and hydrolases have been associated with the degradation of PE, PP or PS, and the molecular mechanisms of substrate recognition and catalysis are mostly unknown.

Recent developments in metagenomics, transcriptomics and comparative genomics have accelerated the finding of novel plastic-associated enzymes from polluted surroundings (Mamtimin et al., 2023; Singh et al., 2026, 2024b). Although most described enzymes are often very substrate-specific, acting mainly on one polymer type or requiring synergistic effects in microbial consortia. In addition, mechanistic understanding of polymer recognition has been largely based on biochemical studies, and few high-resolution structural studies have explained the nature of enzyme interactions with chemically inert carbon–carbon backbone polymers (Shehbas et al., 2026). Therefore, finding multifunctional enzymes capable of degrading several synthetic polymers is one of the primary difficulties in plastic biotechnology.

Hydrolases are among the most varied enzyme superfamilies, catalysing the hydrolysis of ester, amide, peptide and lipid substrates (Chatonnet et al., 2023). Most of the plastic-degrading hydrolases characterised to date are serine hydrolases with cutinases, esterases, lipases and PETases being the classical forms of such enzymes, where catalysis is mediated by the classical Ser-His-Asp catalytic triad (Singh et al., 2024a; Srivastava et al., 2025). However, cysteine-dependent hydrolases in synthetic polymer biodegradation have been comparatively less studied. The enzymes employ a catalytic cysteine residue, whose increased nucleophilicity enables the cleavage of chemically stable substrates under mild physiological circumstances. Cysteine hydrolases of the classical type participate in several biological activities, including proteolysis, ubiquitin processing, and amide hydrolysis, but their possible involvement in the degradation of highly resistant carbon-carbon backbone polymers has been largely unexplored (Pérez-García et al., 2025). Thus, understanding the structural basis of catalysis in cysteine-dependent hydrolases should unveil novel enzymatic pathways that can be exploited to construct broad-spectrum plastic-degrading biocatalysts (Appel and Bertozzi, 2015).

In our previous study, we identified and structurally characterised a novel cysteine-dependent hydrolase, CsdH, from *Rhodococcus opacus* IITR-PK that can degrade polyethylene terephthalate, a heterochain polymer (Aggarwal et al., 2026). In this work, we unravel the molecular mechanism of CsdH-mediated degradation of polypropylene, polyethylene, and polystyrene. Crystal structures of PP, PE, and PS -soaked complexes were determined to highlight substrate-induced conformational changes in the catalytic pocket. Scanning electron microscopy (SEM), Fourier transform infrared spectroscopy (FTIR), thermogravimetric analysis (TGA) and LC–MS/QTOF analyses indicate polymer degradation and degradation products. This work collectively provides the first structural insight into a multifunctional cysteine-dependent hydrolase capable of recognising and degrading PE, PP and PS, thus providing a molecular framework for rational engineering of broad-spectrum enzymes for sustainable biological recycling of mixed plastic waste.

## 2. Materials and methods

### 2.1 Cloning, expression, and purification of CsdH

The cloning, expression, and purification of CsdH hydrolase were performed according to our previously mentioned procedure (Aggarwal et al., 2026).

### 2.2 Degradation study

PP (cat no: 428175), PE (cat no: <u>434272</u>), and PS (cat no: <u>9003-53-6</u>) beads were purchased from Sigma Aldrich (St. Louis, Missouri, USA). To perform degradation study beads were washed with 70 % ethanol and dried for 24 hr at room temperature. and treated in a buffer containing Tris-Cl, pH 8.0, with and without enzyme (1 µM). All samples were incubated at 45 ℃ for 96 hr as optimized in our previous study (Aggarwal et al., 2026). The reaction was quenched by adding 100 % methanol to the reaction mixture, and the precipitation was removed by centrifugation at 14000 x g for 15 min. Then, the beads were washed with 1 % sodium dodecyl sulfate (SDS) in distilled water and 70 % ethanol for SEM, FTIR, TGA, analysis and the supernatant was subjected to LC-MS analysis.

### 2.3 Scanning electron micrograph (SEM) analysis for surface topography changes

The PP, PE, and PS were analyzed to check for any morphological changes or degradation on the surface after treating them with the CsdH enzyme. The air-dried samples were gold-coated (10 nm) using Quorum Q150T S plus and subjected to SEM (FE-SEM, MIRA3 TESCAN, USA) observation.

### 2.4 Attenuated total reflection Fourier transform infrared (ATR-FTIR) spectroscopy analysis

FTIR, a surface-sensitive technique, was used to monitor chemical changes in enzymatically treated PP, PE, and PS beads. Beads were recovered after 96 hr of incubation with CsdH enzyme, thoroughly cleaned in ultrapure distilled water, 1 % SDS, and 75 % ethanol, followed by air drying as per (Srivastava et al., 2024). To obtain infrared spectra, a PerkinElmer spectrometer (Model No. C91158, NJ, USA) and an ATR containing a diamond crystal fixed at an incident angle of 45° were used. The beads were analyzed by transmittance spectra in area between 4000 and 400 cm^-1^, with 120 scans at spectral resolution of 16 cm^-1^ in triplicate using MicroLab software by Agilent Technologies.

### 2.5 TGA analysis

TGA was performed to detect the thermal decomposition of the PP, PE, and PS polymers using a thermogravimetric analyzer (NETZSCH STA 449F3). Approximately 28 mg of each treated bead was used for analysis at 20-800 ℃ with a heating rate of 10 ℃/ min. The experiments were performed in a nitrogen atmosphere with a flow rate of 60 mL/min. Data were analyzed using the Universal Analysis 2000 software.

### 2.6 LC-MS analysis

LC–MS grade methanol and acetonitrile (≥99.9 % purity; J.T Baker, USA) were used throughout. LC–MS-grade formic acid (≥98 % purity; Tokyo Chemical Industry, India) served as the mobile-phase modifier. Type I water suitable for LC–MS was produced using a Klorofil purification system. Analyses were performed on a Waters Xevo G2-XS QT mass spectrometer equipped with an ACQUITY UPLC I-Class system and an electrospray ionization (ESI) source (Waters, USA). Chromatographic separation was performed on an ACQUITY UPLC BEH C18 column (1.7 µm, 2.1 × 50 mm; Waters), operated at a flow rate of 0.35 mL min⁻¹ and an injection volume of 2 μL per sample. The column temperature was maintained at 40 °C, and the autosampler (sample tray) was held at 4 °C. The mobile phases were A, water with 0.1 % (v/v) formic acid, and B, acetonitrile with 0.15 % (v/v) formic acid. The gradient was programmed as follows: 5 % B for 1.0 min; ramp to 100 % B over the next 13.0 min; hold at 100 % B for 4.5 min; return to 5 % B in 1.0 min; re-equilibrate at 5 % B for 0.5 min, giving a total run time of 20.0 min. The QT of source and acquisition parameters were: ESI positive mode, capillary voltage 3.0 kV, cone voltage 40 V, cone gas flow 50 L h⁻¹, desolvation gas flow 600 L h⁻¹, source temperature 120 °C, and desolvation temperature 350 °C. The analyzer was operated in sensitivity mode. Leucine enkephalin was infused as the lock-mass standard for continuous mass-accuracy correction.

### 2.7 Crystallization of CsdH hydrolase

The CsdH protein was concentrated to 15 mg ml^-1^ with the buffer containing 50 mM Tris-HCl and 250 mM NaCl using an Amicon Ultra centrifugal 10 kDa concentrator. Initial crystallization trials were done using various commercially available crystallization screens, including JCSG++ (Jena Bioscience) and Wizard I and II (Rigaku). The crystallization drop includes 1 µl of protein mixed with 1 µl of reservoir solution, equilibrated with 60 µl of reservoir solution. The optimal condition for obtaining diffraction data was found to be 0.2 M sodium fluoride, 20 % PEG 3350, with a protein to reservoir ratio of 1:1. The crystals were cryoprotected in a solution containing 30 % ethylene glycol and mother liquor solution, followed by flash freezing in liquid nitrogen at a temperature of 100 K. To elucidate the protein’s catalytic mechanism, it was soaked with PP, PE, PS, and styrene, each at a concentration of 10 mM, for 10 minutes (Mahto et al., 2025, 2022).

### 2.8 X-ray data collection and structure determination

The X-ray diffraction data were collected at the in-house facility of the Macromolecular Crystallographic Unit, Institute Instrumentation Centre, Indian Institute of Technology Roorkee (IIT Roorkee), India. Detailed statistics for data collection and processing are summarized in **Table 1**. The data quality was evaluated using Aimless of the CCP4i suite. The CsdH-apo (PDB ID 26MZ) was used as a search model for molecular replacement using CCP4i. Model building was performed using COOT, iterative model building, electron density maps, and REFMAC5 for manual fitting into the model, both integrated into the CCP4i2 suite. Non-crystallographic symmetry (NCS) restraints were applied throughout the hydrolase data refinement process. Water molecules were incorporated during the final stages of structure refinement, where the *F*o-*F*c map showed values above 3σ. All figures of protein and ligand structures were prepared using PyMol (Delano, 2002). Polder maps were constructed using PHENIX. The active site cavity analysis was done using the MOLE online server (Raček et al., 2025). The tunnel analysis was done using CAVER 3.0.3 (Chovancova et al., 2012).

**Table 1.** Data processing and refinement statistics for CsdH complex structures.

| Parameter | CsdH-Polypropylene<br>(26RX) | CsdH-Polyethylene<br>(44FL) | CsdH-Polystyrene<br>(26RM) | CsdH-Styrene<br>(26RB) |
| --- | --- | --- | --- | --- |
| Resolution range | 28.02- 3.0<br>(3.107 - 3.0)<sup>a</sup> | 28.33- 2.9<br>(3.004 - 2.9)<sup>a</sup> | 27.67-2.6<br>(2.693-2.6)<sup>a</sup> | 28.59- 2.5<br>(2.589- 2.5)<sup>a</sup> |
| Space group | *C* 2 2 2<sub>1</sub> | *C* 2 2 2<sub>1</sub> | *C* 2 2 2<sub>1</sub> | *C* 2 2 2<sub>1</sub> |
| Unit cell<br>*a b c* (Å) | 61.40<br>136.90<br>187.80 | 62.28<br>136.51<br>188.38 | 62.18<br>136.92<br>187.92 | 62.27<br>137.10<br>188.76 |
| $\alpha \beta \gamma$ (°) | 90.00 90.00 90.00 | 90.00 90.00 90.00 | 90.00 90.00 90.00 | 90.00 90.00 90.00 |
| Multiplicity | 13.0 (13.6) | 6.5 (6.6) | 6.0 (6.3) | 14.0 (8.5) |
| Completeness (%) | 99.43 (100.00) | 99.29 (99.27) | 99.48 (99.84) | 99.75 (99.75) |
| I/σ (I) | 21.75 (5.50) | 22.19 (5.43) | 16.35 (4.95) | 36.74 (4.97) |
| Wilson B-factor (Å<sup>2</sup>) | 31.20 | 32.44 | 31.46 | 35.23 |
| R<sub>merge</sub><sup>b</sup> | 0.139(0.458) | 0.101(0.336) | 0.122(0.405) | 0.084(0.624) |
| R<sub>pim</sub> | 0.040(0.128) | 0.044(0.141) | 0.052(0.170) | 0.023(0.234) |
| CC1/2 | 0.995(0.921) | 0.994(0.931) | 0.993 (0.908) | 0.997(0.505) |
| Reflections used in refinement | 16195(1600) | 18123(1773) | 25018 (2478) | 28380(2800) |
| Reflections used for R-free | 822(67) | 906(94) | 1218(112) | 1427(135) |
| R<sub>work</sub><sup>c</sup> | 0.204(0.206) | 0.201(0.248) | 0.1955(0.218) | 0.196(0.424) |
| R<sub>free</sub> | 0.263(0.283) | 0.250(0.312) | 0.238(0.273) | 0.226(0.498) |
| Number of non-hydrogen atoms | 6590 | 6595 | 6641 | 6907 |
| Macromolecules | 6543 | 6551 | 6548 | 6559 |
| Ligands | 24 | 14 | 36 | 193 |
| Solvent | 23 | 30 | 57 | 155 |
| Protein residues | 864 | 865 | 866 | 887 |
| RMS (bonds) (Å)<sup>d</sup> | 0.008 | 0.009 | 0.009 | 0.022 |
| RMS (angles) (degrees) | 1.57 | 1.72 | 1.74 | 1.88 |
| Favored (%) | 95.54 | 97.10 | 99.06 | 97.19 |
| Allowed (%) | 3.87 | 2.58 | 0.82 | 1.99 |
| Outliers (%) | 0.59 | 0.32 | 0.12 | 0.82 |
| Rotamer outliers (%) | 3.94 | 4.12 | 4.38 | 4.25 |
| Clash score | 4.17 | 4.69 | 4.40 | 4.52 |
| Average B-factor (Å<sup>2</sup>) | 31.61 | 33.56 | 32.36 | 34.32 |
| Macromolecules (Å<sup>2</sup>) | 31.71 | 34.56 | 32.52 | 30.40 |
| Ligands | 35.03 | 34.18 | 33.12 | 35.51 |
| Solvent (Å<sup>2</sup>) | 36.46 | 38.12 | 38.64 | 37.21 |

**a** Values in parentheses are for the highest resolution shell.
**b** $R_{\text{merge}} = \sum |I - \langle I \rangle| / \sum I$
**c** $R = \sum |F_{\text{obs}}| - |F_{\text{calc}}| / \sum |F_{\text{obs}}|$ . The $R_{\text{free}}$ is the R calculated on the 5 % reflections excluded for refinement
**d** RMS is the root mean square

## 3. Results

### 3.1 Weight loss assay of enzymatically degraded polymers

The enzymatic degradation of various polymers, PP, PE, and PS, was evaluated through weight loss measurements following treatment **(Figure 1A)**. PE showed the maximum degradation with a mean weight loss of 12.14 ± 0.88 %, indicating the highest degradation among the tested polymers. PP exhibited degradation at an intermediate level with a mean weight loss of 6.37 ± 1.23%, and PS showed the lowest degradation with a mean weight loss of 5.87 ± 1.00 %. These results establish the differential enzymatic degradability of polymer types, with PE being the most affected under the tested conditions. These morphological changes were further validated by SEM observations. The structures of 3-PS,3-PE, and 3-PP are shown **(Figure 1B)**.

**Figure 1:**
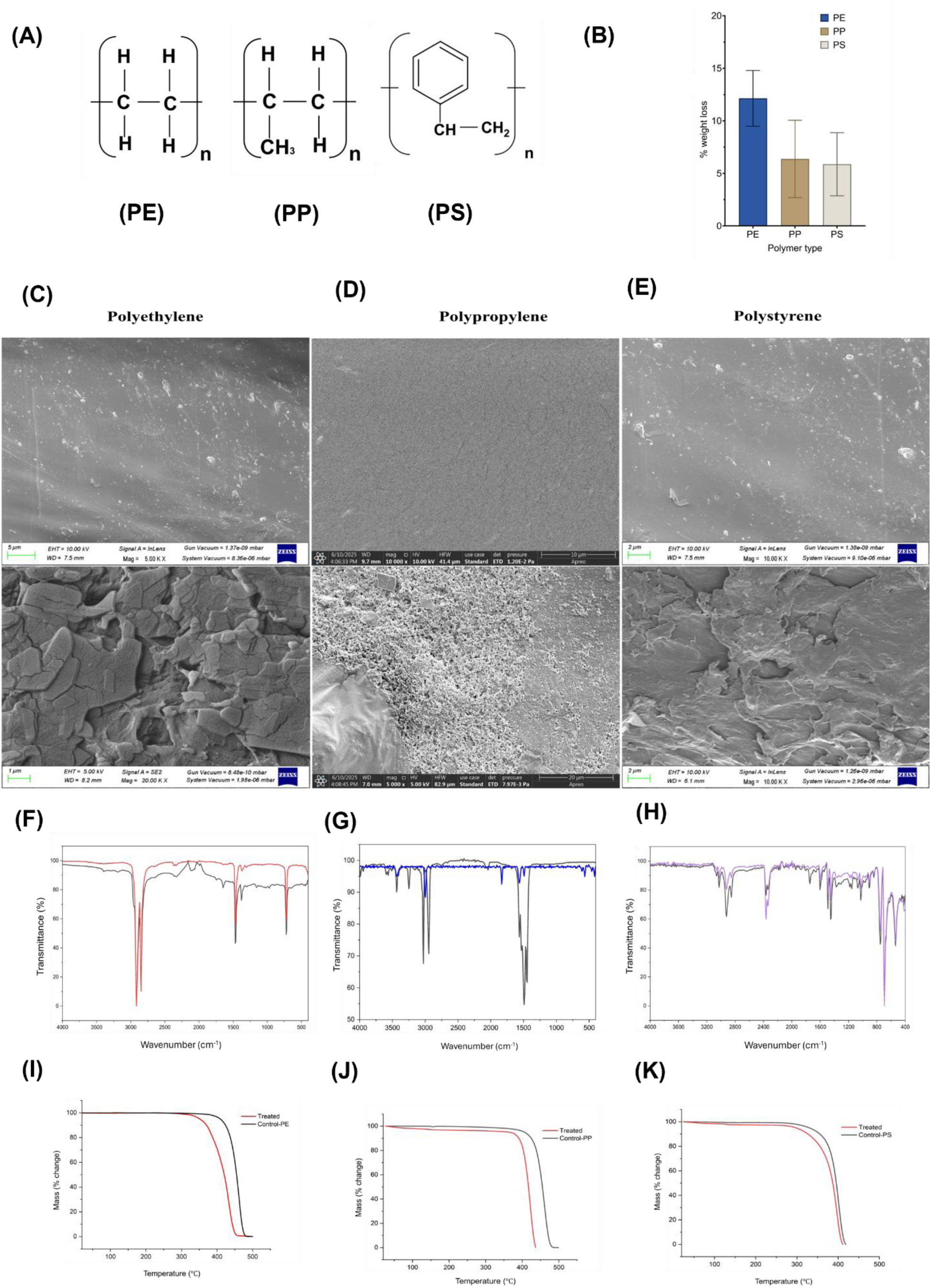
**(A)** Structure of PE, PP and PS. **(B)** Weight loss evaluation of polyethylene, polypropylene, and polystyrene. Scanning electron micrograph of **(C)** control (black) and treated PE (red), **(D)** control (black) and treated PP (blue), **(E)** control (black) and treated PS (purple). FTIR analysis of **(F)** PE with carbonyl index increased by ∼7.8 %, **(G)** PP with reduced CI value of 0.9966, **(H)** PS with aromatic C–H stretching peak near 3051 cm⁻¹ exhibited a slight increase (+0.96 %) after treatment. Thermogravimetric analysis of **(I)** PE with degradation was observed at 409.0 °C, and 50 % weight loss (T_50_) occurred at 454.9 °C, **(J)** PP with degradation at 368 °C and reached 50 % weight loss (T_50_) at 399.9 °C, and **(K)** PS with degradation at 330.2 °C, and the temperature corresponding to 50% weight loss (T_50_) was 395.5 °C.

### 3.2 SEM analysis

The untreated PE showed a compact, relatively rough texture, characteristic of semicrystalline polymers. After enzymatic treatment, the PE surface exhibited increased roughness, micro-cracking, and granular fragmentation, especially visible at higher magnifications (scale bar = 1-5 µm). The observed textural change suggests partial enzymatic modification at the polymer surface, although complete degradation was not seen. This moderate alteration implies limited enzymatic interaction with the hydrophobic PE surface **(Figure 1C)**.

The untreated PP displayed a smooth morphology typical of this hydrophobic polymer. After enzymatic exposure, significant surface irregularities and shallow depressions appeared, as seen at higher magnification (scale bar: 10-20 µm). The limited morphological change indicates that PP is relatively resistant to enzymatic degradation, likely due to its non-polar backbone and lack of hydrolyzable functional groups **(Figure 1D).**

Similarly, after enzymatic treatment, the PS surface developed visible grooves, fissures, and minor delamination features (scale bar =2 µm). These morphological changes suggest that enzymatic exposure led to partial surface etching, possibly through oxidative degradation mechanisms **(Figure 1E)**.

### 3.3 FTIR analysis

Enzyme-treated samples of polyethylene (PE) showed significant spectral changes when compared to the control group. The carbonyl index experienced an increase of approximately 7.8%, indicating the oxidation process and the incorporation of C=O groups into the polymer matrix. The alkene region (700-900 cm⁻¹) exhibited an increase of approximately 26.8 %, indicating the occurrence of chain scission and unsaturation. Furthermore, the C-O stretching band (1050-1150 cm⁻¹) showed an increase of approximately 28.7 %, which aligns with the formation of oxygenated species such as alcohols and esters during the process of enzymatic degradation. These findings validate the oxidative and hydrolytic degradation of polyethylene chains **(Figure 1F)**.

Polypropylene’s (PP) FTIR spectra revealed clear signs of enzymatic degradation, characterised by subtle yet significant changes in the distinctive absorption bands. The production of oxygen-containing functional groups during enzymatic oxidation is reflected by a slight increase in absorbance in the 1700–1800 cm⁻¹ range, which is associated with C=O stretching vibrations. At the same time, a decrease in the intensity of the 2800–3000 cm⁻¹ region indicates potential partial breakage of the polypropylene backbone. This change is linked to the symmetric and asymmetric stretching vibrations of CH_2_ and CH_3_ groups. Alterations in the molecular structure and local crystallinity of the polymer were observed in the fingerprint region (<1500 cm⁻¹). Together, these spectral changes indicate that polypropylene experienced oxidative surface degradation and initial chain scission as a result of enzymatic treatment. This process modified the polymer’s structure while primarily preserving the integrity of its backbone **(Figure 1G)**.

FTIR analysis of polystyrene (PS) showed only minor spectral differences between the untreated and enzyme-treated samples, suggesting that there was limited modification of the polymer. A slight increase in the O–H stretching bands at approximately 3653 and 3448 cm⁻¹ was noted in the treated samples, indicating the formation of a small quantity of hydroxyl-containing oxidation products on the polymer surface. The aromatic C–H stretching vibration at 3051 cm⁻¹ showed a slight increase of approximately 0.96 %, which does not suggest any substantial change in the structure of the aromatic backbone. In a similar manner, the aliphatic C–H stretching band at 2881 cm⁻¹, the carbonyl region around 1757 cm⁻¹, the aromatic skeletal vibration at 1480 cm⁻¹, and the out-of-plane aromatic C–H bending vibration at 726 cm⁻¹ exhibited minimal to no significant change after enzyme treatment. Overall, the FTIR results suggest that the enzymatic treatment led to only minor surface oxidation, while maintaining the integrity of the characteristic aromatic structure of polystyrene **(Figure 1H)**.

### 3.4 TGA analysis

Thermogravimetric analysis (TGA) of polypropylene (PP), polystyrene (PS), and polyethylene (PE) samples was performed under a nitrogen atmosphere to evaluate the effect of enzymatic treatment on their thermal stability. All control samples exhibited characteristic single-step degradation profiles typical of their polymer backbones, whereas enzyme-treated samples consistently showed a shift of degradation temperatures to lower values, indicating reduced thermal stability.

For polyethylene (PE) samples, control PE showed thermal decomposition in the range of 409-454 °C. The onset of degradation for the control sample was observed at 409.0 °C, and 50 % weight loss (T50) occurred at 454.9 °C. Only trace amounts of residue (0.0009 %) remained at the end of the heating cycle. In contrast, the enzymatically treated PE sample exhibited an earlier onset of degradation at 351.4 °C and reached 50 % weight loss at 420.0 °C. The observed decrease in T_onset_ and T_50_ values suggests that the enzymatic treatment has notably diminished the thermal stability of polyethylene (PE). This decrease in stability aligns with enzymatic modifications that result in polymer chain cleavage, a reduction in molecular weight, and changes in crystallinity **(Figure 1-I)**. Enzymatic treatment resulted in a shift of both T_onset_ and T_50_ values to lower temperatures across all tested polymers, indicating a reduction in thermal stability. This reduction in stability can be linked to the enzymatic cleavage of polymer chains, a decrease in molecular weight, and changes in crystallinity. The observed changes align with previous findings regarding the microbial and enzymatic degradation of synthetic polymers **(Figure 1-I)**.

The weight loss in the control polypropylene sample occurred in the temperature range of 368-399 °C and showed an onset of degradation at 368 °C and reached 50 % weight loss (T_50_) at 399.9 °C. In comparison, the enzymatically treated PP sample exhibited an earlier onset of degradation at 349.3 °C and reached 50 % weight loss at 397.2 °C **(Figure 1J)**. This reduction in T_onset_ and T_50_ values indicates a decrease in the thermal stability of the PP after enzymatic attack. The results clearly show that the degradation temperature shifted to the lower side after enzymatic treatment, revealing a loss of thermal stability.

Similarly, polystyrene (PS) samples exhibited thermal decomposition in the temperature range of 330-395 °C. and showed onset of degradation at 330.2 °C, and the temperature corresponding to 50 % weight loss (T_50_) was 395.5 °C. The enzymatically treated PS sample showed an earlier onset of degradation at 298.0 °C and reached 50 % weight loss at 386.9 °C. The observed shift of degradation temperatures to the lower side demonstrates polymer chain scission and reduction in molecular weight, which weakens the structural integrity of polystyrene **(Figure 1K)**.

### 3.5 LC-MS analysis for detection of degradation metabolites

In order to assess the enzymatic degradation capabilities of the hydrolase on synthetic polymers, polypropylene (PP), polyethylene (PE), and polystyrene (PS) beads were separately incubated with the enzyme under carefully optimised conditions. The supernatants obtained were analysed using LC-MS/Q-TOF to identify significant degradation metabolites **(Supplementary Table S1)**.

In the case of PE, characteristic aliphatic and oxidative degradation products were observed, including hexane (m/z 125.07) **(Figure 2A)**, acetic acid (m/z 98.98), and pentane (m/z 95.08) **(Supplementary Figure S1),** suggesting terminal oxidation and chain shortening through β-oxidation-like mechanisms.

**Figure 2:**
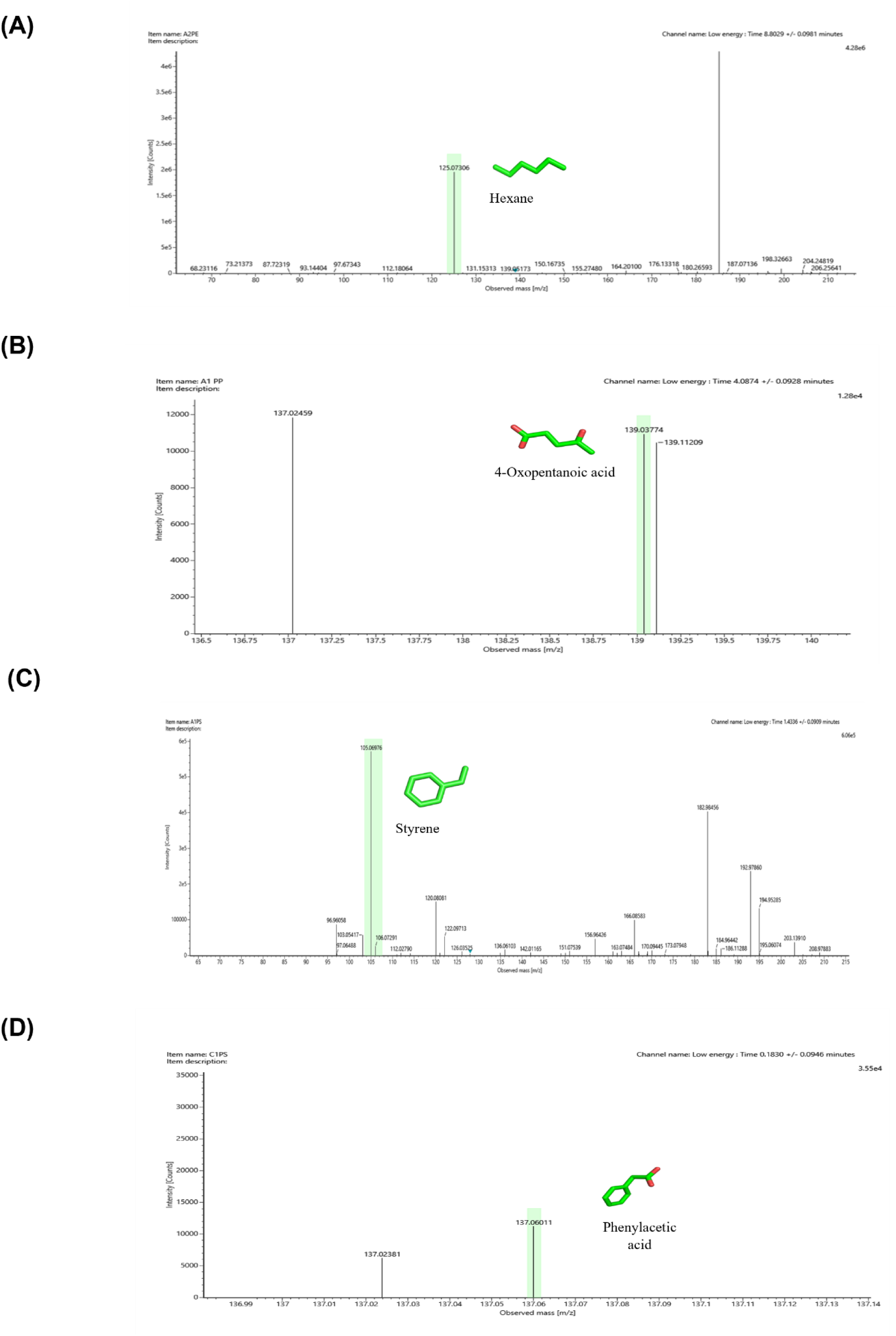
Liquid chromatography–mass spectrometry (LC–MS) analysis of the major PE, PP, and PS degradation products generated during enzymatic hydrolysis by CsdH. Representative mass spectra are shown for **(A)** Hexane (m/z 125.07) ([M+ K]⁺), **(B)** 4-oxopentanoic acid (m/z 139.03) ([M+ Na]⁺), **(C)** Styrene (m/z 105.0699) ([M+ H]⁺), and **(D)** Phenylacetic acid (m/z 137.0693).

For PP, the formation of 4-oxopentanoic acid (m/z 139.03) **(Figure 2B),** 2-butenoic acid (m/z 87.04), and acetic acid (m/z 98.98) (**Supplementary Figure S2)** were identified. These metabolites are indicative of beta-oxidation-linkage breakdown of the hydrocarbon chains and confirm the partial oxidation and fragmentation of PP chains into carboxylic acids and ketonic intermediates.

For PS, preliminary metabolite analysis revealed a diverse set of aromatic degradation products, confirming the oxidative and hydrolytic cleavage of the PS backbone. Major identified compounds included styrene (m/z 105.0699) **(Figure 2C)** and styrene oxide (m/z 121.0666), indicating initial oxidation of the vinyl group. Further oxidation and ring-cleaving reactions led to the formation of benzoic acid (m/z 123.0431), phenylacetic acid (m/z 137.0693) **(Figure 2D)**, p-hydroxybenzoic acid (m/z 139.0399), and *p*-hydroxyphenylacetic acid (m/z 153.0541) (**Supplementary Figure S3)**. The presence of such structurally diverse low molecular weight products suggests that the hydrolase catalyzes both initial depolymerization and subsequent oxidation steps.

The detection of both polymer-specific and common degradation products across all three plastics reflects the broad substrate tolerance and catalytic versatility of the hydrolase. These findings validate the multi-substrate degradation capability of the enzyme and its potential application in biotechnological strategies for mixed plastic waste bioremediation.

### 3.6 Structural basis and functional validation of CsdH-mediated homopolymer depolymerization

#### 3.6.1 CsdH-Polyethylene complex structure and its biodegradation

CsdH-PE structure determined at 2.9 Å resolution, having R_work_ and R_free_ values of 20.1 % and 25.0 % **(Table 1),** respectively, in space group *C*222_1,_ revealed that PE undergoes complete degradation within the crystal and forms a hexane molecule rather than the intact substrate as shown in the polder map contoured at 3σ near the active site triad (Cys160-Lys127-Asp39) **(Figure 3A-B)**. Thus, in crystallo, turnover demonstrates that CsdH is catalytically active under crystallization conditions. The structure of CsdH reveals how it binds to breakdown products of polyethylene. The active site is defined by a network of catalytic residues Cys160, Lys127 and Asp39, and a largely hydrophobic substrate-binding pocket is formed by Trp93, Phe44, Asn100, Ala130 and Thr156 with contributions from Arg185. Notably, a hexane molecule was detected in the catalytic cavity, suggesting that the active site can accommodate a short aliphatic hydrocarbon chain. The proximity of the hydrocarbon to the catalytic Cys160 indicates that the hydrophobic pocket may act as a substrate-recruitment and positioning site for aliphatic fragments generated during polyethylene oxidation.

**Figure 3:**
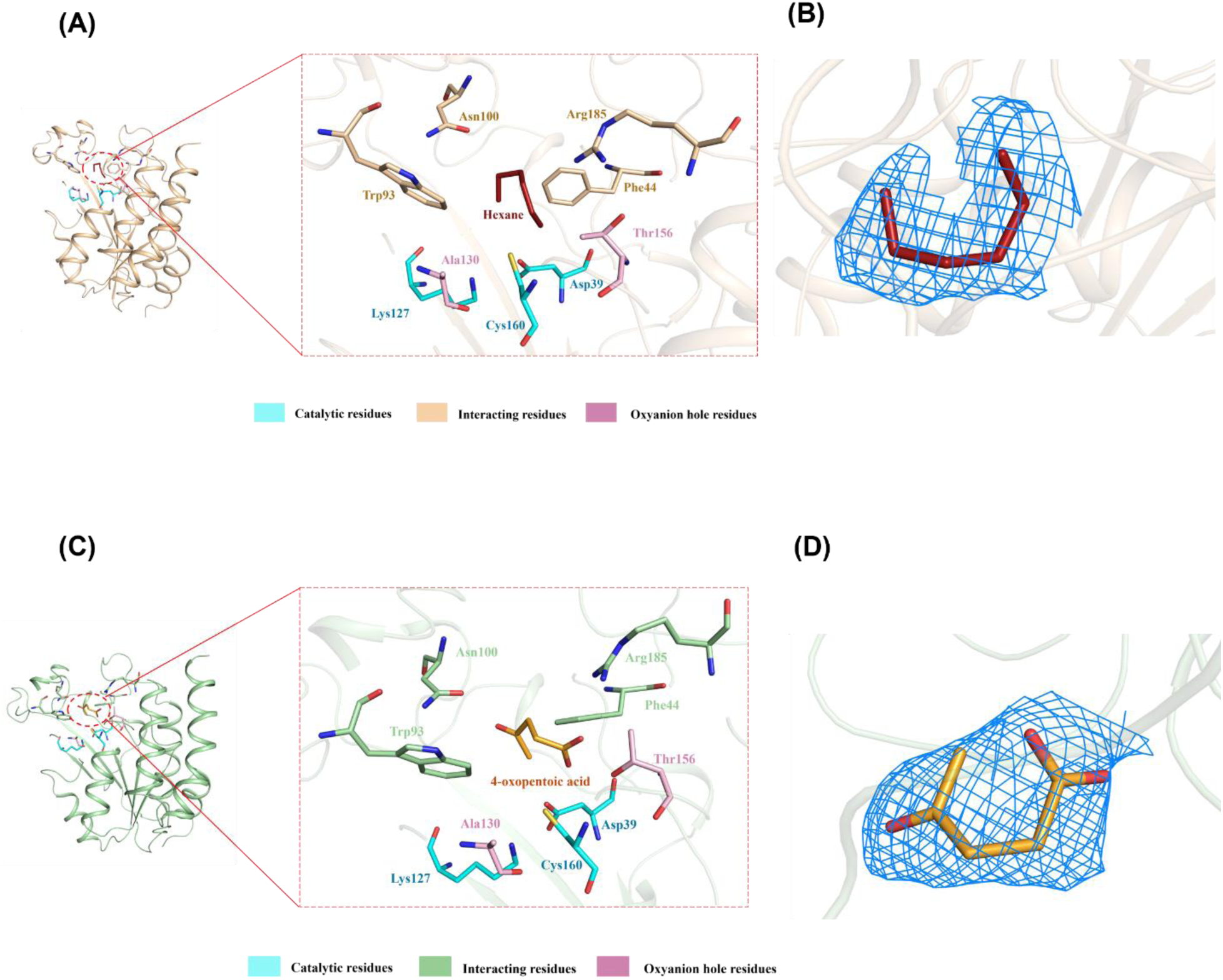
**(A)** Active-site interactions of hexane, the hydrolysis product of polyethylene, following ligand soaking experiments. The ligand (red sticks) is anchored through hydrogen-bond interactions with Trp93, Ala130, Phe44, Thr156, Arg185, and Asn100. The catalytic triad residues Cys160–Lys127–Asp39 are positioned proximal to the ligand-binding site. **(B)** Polder map contoured at 3σ showing the electron density of hexane, the hydrolysis product of polyethylene. **(C)** Active-site interactions of 4-oxopentanoic acid, the hydrolysis product of polypropylene, following ligand soaking experiments. The ligand (mustard-coloured sticks) is anchored via hydrogen-bond interactions with Trp93, Ala130, Phe44, Thr156, Arg185, and Asn100. The catalytic triad residues Cys160–Lys127–Asp39 are positioned proximal to the ligand-binding site. **(D)** Polder map contoured at 3σ showing the electron density of 4-oxopentanoic acid, the hydrolysis product of polypropylene.

It is unlikely that CsdH directly cleaves the unmodified PE C–C backbone, as hydrolyzable ester bonds are absent in native polyethylene. Instead, we suggest that CsdH acts on chemically oxidized and shorter chain PE-derived intermediates formed in the early stages of polyethylene oxidation. In this model hydrophobic fragments derived from PE are recruited into the active-site cavity where the catalytic network of Asp39, Lys127 and Cys160 activates the cysteine nucleophile. Cys160 then attacks an electrophilic carbonyl group in an oxidized PE-derived substrate generating a tetrahedral intermediate. This high energy intermediate is proposed to be stabilized by the surrounding oxyanion stabilizing region involving the backbone amide groups of Thr156 and Ala130. The intermediate then collapses and Cys160 is regenerated by hydrolysis, cleaving the activated substrate and yielding shorter chain products.

The LC–MS detection of hexane, pentane, butenoic acid and acetic acid is in agreement with progressive fragmentation and oxidation of polyethylene-derived carbon chains. In particular, the detection of hexane in the crystal structure of CsdH and its identification as one of the degradation products further supports the notion that the enzyme can recognize short aliphatic chains. However, the crystallographically observed hexane should be taken as an indication that the substrate pocket is occupied rather than as a conclusive proof that hexane is directly produced by the catalytic reaction of CsdH. The structural and LC–MS data are consistent with a model in which CsdH contributes to the downstream processing of oxidized PE fragments via cysteine-dependent hydrolysis and allows the conversion of larger oxygenated polymer-derived intermediates to smaller aliphatic and acidic products.

#### 3.6.2 Polypropylene degradation at the active site

In the polypropylene-soaked state, the crystal structure of the complex was determined at 3.0 Å, with R_work_ and R_free_ values of 20.4 % and 26.3 %, respectively **(Table 1).** In the CsdH–PP complex, polder map contoured at 3σ reveals clear electron density for the product, 4-oxopentanoic acid (laevulinic acid) (LA), occupying the active-site cavity **(Figure 3C)**, and is further stabilized by interactions with Trp93, Ala130, Phe44, Thr156, Arg185 and Asn100 indicating that PP chain scission occurred prior to or during crystallization and that the structure captures a post-catalytic product-bound state **(Figure 3D)**. This binding mode suggests that CsdH has a substrate-recognition cavity to bind both hydrophobic and oxygenated compounds derived from polypropylene. Asn100, Arg185 and Thr156 provide additional polar contacts that help to orient the substrate/product, while the hydrophobic residues Trp93 and Phe44 are probably involved in the recognition and packing of the aliphatic part of the substrate. The carbonyl oxygen of laevulinic acid is a hydrogen-bond acceptor and is stabilised by hydrogen bonds from the backbone amide groups of Thr156 and Ala130, thus defining an oxyanion-stabilising environment within the active site. This is particularly noteworthy. This structural arrangement is in agreement with the proposed role of these residues in the stabilization of a negatively charged oxygen during catalysis.

Also, apo and PP-bound structures were compared to reveal the rearrangement of the catalytic network associated with the ligand. The distance between Cys160 Sγ and Asp39 O increased from 3.35 Å in apo structure to 3.59 Å in the PP bound state and the distance between Cys160 Sγ and Lys127 Nε increased from 3.40 to 3.57 Å. In addition, the Asp39–Lys127 interaction was shifted by approximately 0.45 Å in position. These modifications indicate that the catalytic residues undergo local rearrangements upon binding the PP-derived ligand/product, which is consistent with an active-site adaptation occurring during substrate recognition and catalysis. However, with the present resolution, these geometrical changes can best be interpreted as structural evidence of rearrangement of the catalytic network, rather than as direct evidence for specific changes in protonation state. From the structural observations, we propose that CsdH is involved in the downstream degradation of oxidatively functionalized polypropylene-derived intermediates and does not directly cleave the chemically inert C–C backbone of pristine PP. In the proposed mechanism, the initial oxidation of PP yields shorter-chain oxygenated intermediates with electrophilic carbonyl functionalities. These compounds are drawn into the CsdH active-site cavity by hydrophobic and polar interactions. The Cys160 thiol is activated by the Cys160–Lys127–Asp39 catalytic network to nucleophilic attack an electrophilic carbonyl center of the oxygenated PP-derived substrate. The backbone amide groups of Thr156 and Ala130 are expected to stabilize a negative charge on the oxygen that is formed when the tetrahedral intermediate is created. The intermediate would then collapse to produce a covalent acyl-enzyme intermediate. Hydrolytic regeneration of CsdH and release of smaller degradation products such as laevulinic acid would then occur.

#### 3.6.3 Polystyrene degradation at the active site

Similarly, we soaked the crystal with the substrate polystyrene, which provides the conformational response of the active site to ligand binding. The CsdH–PS structure was determined to 2.6 Å resolution with R_work_ 19.5 % and R_free_ 23.8 % in space group *C*222_1_ **(Table 1).** Polder map contoured at 3σ reveals clear density for the styrene **(Figure 4D)** and phenylacetic acid **(Figure 4B)** as the product near the catalytic triad (Cys160-Lys127-Asp39), which was also suggested by LC-MS analysis, indicating that enzymatic hydrolysis occurred during crystallization. The C1 carbon of phenylacetic acid **(Figure 4C)** is positioned 2.9 Å from Cys160 Sγ, slightly further than the 2.69 Å observed for styrene **(Figure 4A)**, consistent with the lower catalytic efficiency of CsdH toward styrene relative to its natural polymer (PS) **(Figure 4D)**. The negatively charged oxygen is stabilized by the oxyanion hole formed by Thr156 and Ala130, releasing phenylacetic acid as the degradation product.

**Figure 4:**
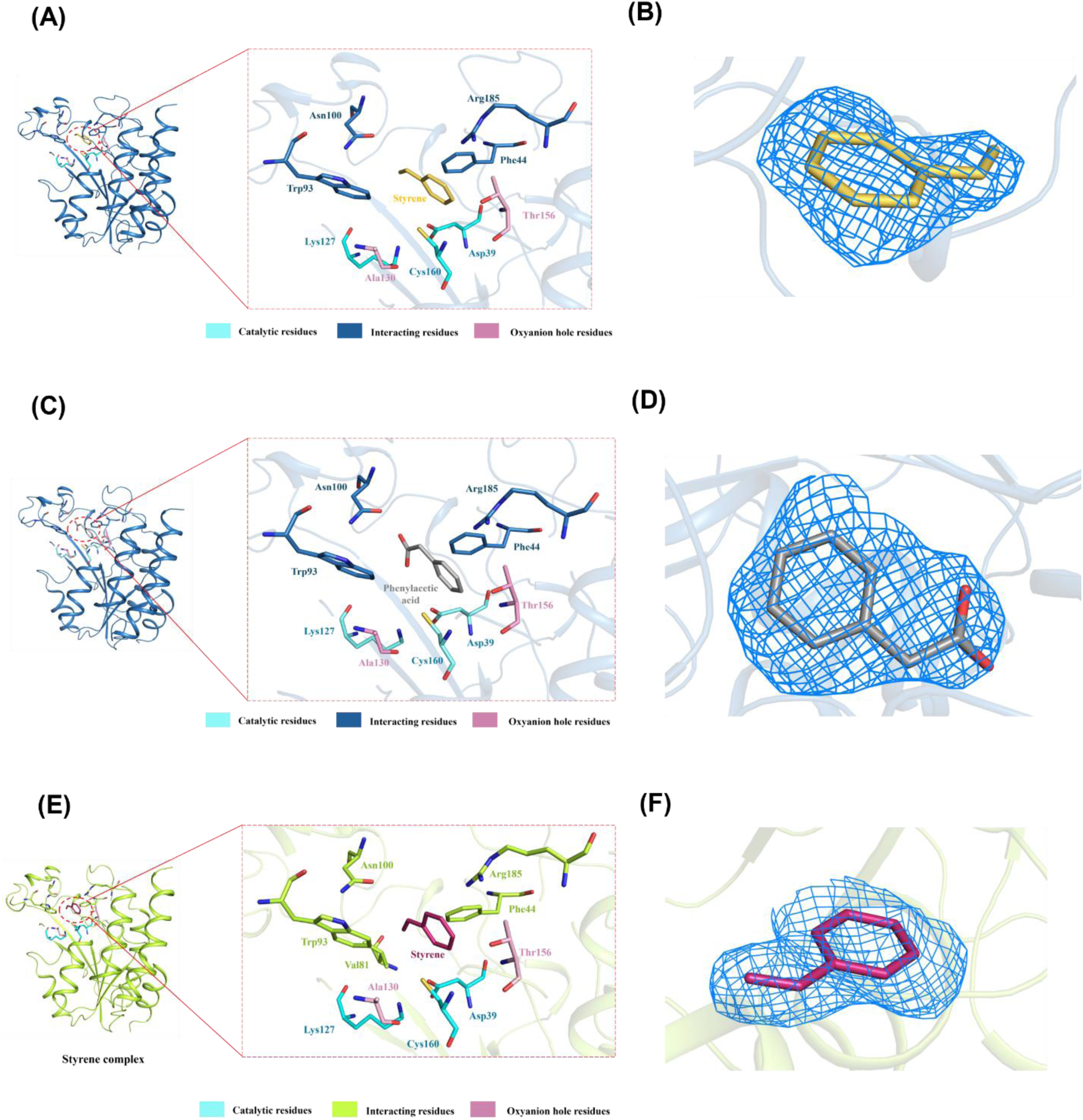
**(A)** Active-site interactions of styrene, the hydrolysis product of polystyrene, following ligand soaking experiments. The ligand (golden colour sticks) is anchored through hydrogen-bond interactions with Thr156, Arg185, and Asn100, while aromatic stacking and hydrophobic interactions with Phe44 contribute to substrate recognition. The catalytic triad residues Cys160–Lys127–Asp39 are positioned proximal to the ligand-binding site. **(B)** Polder map contoured at 3σ showing the electron density of styrene, the hydrolysis product of polystyrene. **(C)** Active-site interactions of phenylacetic acid, the hydrolysis product of polystyrene, following ligand soaking experiments. The ligand (grey colour sticks) is anchored through hydrogen-bond interactions with Thr156, Arg185, and Asn100, while aromatic stacking and hydrophobic interactions with Phe44 contribute to substrate recognition. The catalytic triad residues Cys160–Lys127–Asp39 are positioned proximal to the ligand-binding site. **(D)** Polder map contoured at 3σ showing the electron density of phenylacetic acid. **(E)** Active-site interactions of styrene, following ligand soaking experiments. The ligand (pink colour sticks) is anchored through van der Waals interactions with Thr156, Arg185, and Asn100, while aromatic stacking and hydrophobic interactions with Phe44 contribute to substrate recognition. The catalytic triad residues Cys160–Lys127–Asp39 are positioned proximal to the ligand-binding site. **(F)** Polder map contoured at 3σ showing the electron density of styrene.

Strikingly, comparison of the apo and polystyrene complex structures reveals an 11° angular deviation in the aromatic side chain of Phe44 upon product binding. This conformational rotation repositions the Phe44 phenyl ring to form a tighter hydrophobic enclosure around the phenylacetic acid aromatic ring, consistent with an induced-fit gating mechanism in which Phe44 acts as a conformational latch that closes over the substrate upon active site engagement.

### 3.6.4 CsdH–Styrene complex structure

To further confirm the binding pocket, we soaked the crystal in styrene and found styrene at the catalytic site, at a resolution of 2.5 Å, with R_work_ 19.6 % and R_free_ 22.6 % in space group *C*222*_1_* **(Table 1)**. The C1 carbon of styrene is positioned 2.69 Å from the Sγ of Cys160. Polder map contoured at 3σ reveals clear density for the styrene **(Figure 4F)** and is further stabilized by van der Waals contacts with Val81, Ile197, Asn100, Arg185, Phe44, and Trp93, a binding network that partially overlaps with the phenylacetic acid-binding contacts in the CsdH-PS complex **(Figure 4E)**, suggesting that both aromatic products compete for the same hydrophobic binding cavity during sequential depolymerization. The close approach of styrene C1 to Cys160 Sγ is also consistent with styrene’s role as a mild competitive inhibitor at high substrate concentrations.

#### 3.6.5 Cavity analysis of apo-CsdH and complex with homopolymers

Cavity analysis revealed substantial substrate-dependent remodelling of the active-site pocket, despite minimal overall architectural changes. The calculated monomer volumes remained relatively conserved among the apo and substrate-bound structures, ranging from 16,308 to 16,843 Å³, indicating that substrate accommodation does not involve major expansion or contraction of the overall protein structure. In contrast, the active-site region displayed pronounced changes in volume, showing an increase from 681 Å³ in the apo structure to 923 Å³, 1,013 Å³, 1,392 Å³, and 1,413 Å³ in the styrene, polystyrene, polypropylene, and polyethylene complexes, respectively. Thus, the cavity expanded by approximately 35.5 % for styrene, 48.8 % for polystyrene **(Figure 5A-E)**.

**Figure 5:**
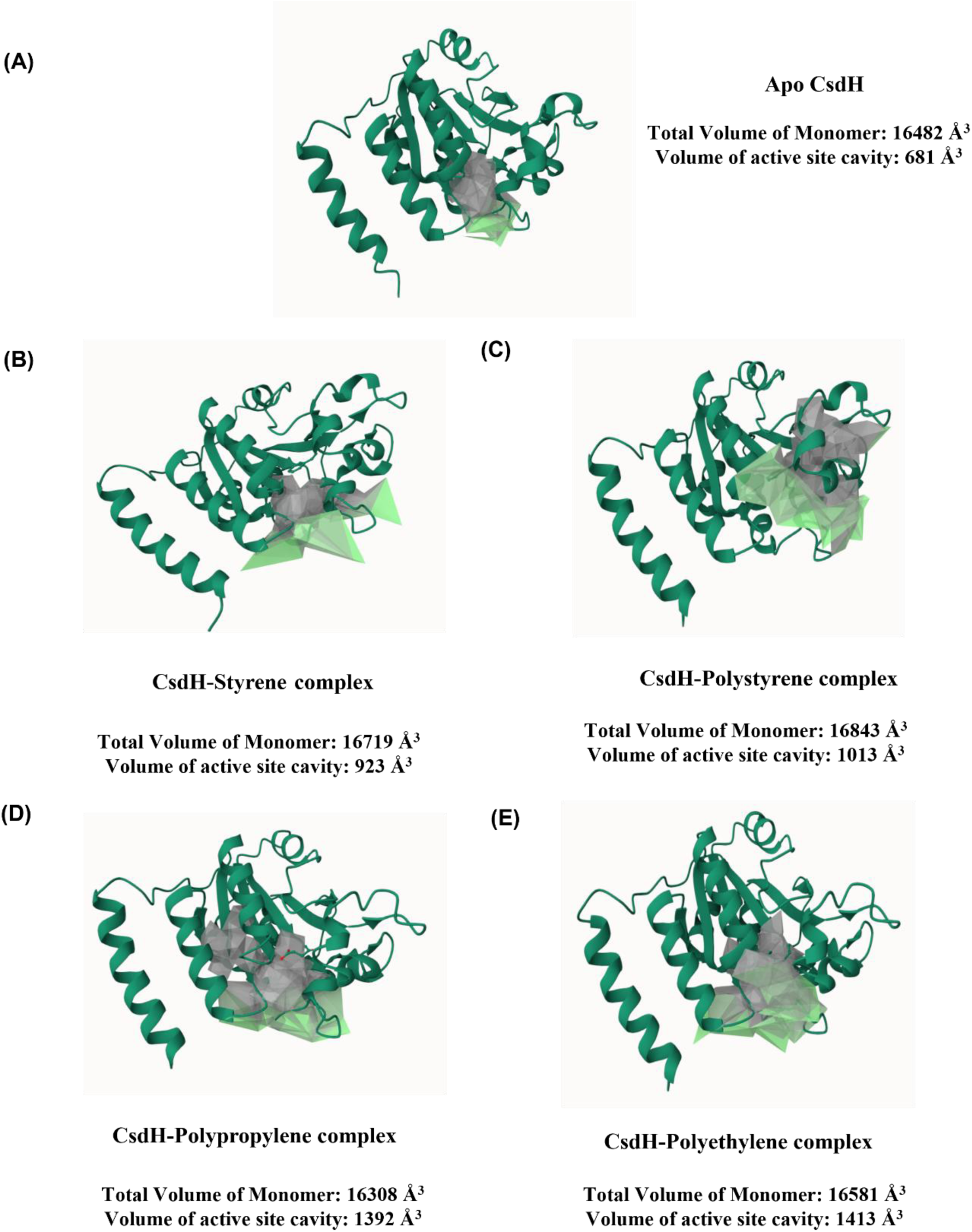
Cavity analysis of (A) apo-CsdH and complex with (B) styrene, (C) polystyrene, (D) polypropylene, and (E) polyethylene. The active-site region displayed pronounced changes in volume, showing an increase from 681 Å³ in the apo structure to 923 Å³, 1,013 Å³, 1,392 Å³, and 1,413 Å³ in the styrene, polystyrene, polypropylene, and polyethylene complexes, respectively. Green and grey colours indicate the different views of the protein and the active site volume

These observations suggest that the protein might possess an adaptable active-site architecture capable of undergoing localized structural rearrangements to accommodate substrates of varying sizes and shapes **(Supplementary Table S2)**. Tunnel analysis revealed one major and another minor access tunnel in both the apo and substrate-bound crystal structures, with Tunnel 1 consistently representing the preferred major pathway owing to its higher throughput (0.88 -0.93) and lower cost (0.07 -0.12) compared with Tunnel 2 (throughput 0.43 -0.53 and cost 0.61 -0.82).

Substrate binding resulted in moderate changes in tunnel dimensions, particularly the bottleneck radius and length, while the overall tunnel architecture remained conserved. Tunnel 1 exhibited bottleneck radii of 1.35-2.27 Å across the substrate complexes, with the polypropylene complex showing the widest bottleneck (2.27 Å) and the shortest radius (1.41 Å), indicative of a highly accessible pathway **(Supplementary Table S3)**.

In contrast, Tunnel 2 remained narrower and longer. The relatively small variation in tunnel curvature suggests that it may not be a preferred choice of substrate entry or exit. Collectively, these results indicate that the protein retains a conserved principal access tunnel, which is tunnel 1, while undergoing subtle substrate-dependent remodelling to accommodate molecules of different sizes and shapes **(Supplementary Figure S4)**.

## 4. Discussion

This study shows that CsdH is a promising multifunctional cysteine-dependent hydrolase that can interact with and degrade three of the most persistent synthetic polymers in the environment, polyethylene (PE), polypropylene (PP) and polystyrene (PS). Compared to most of the plastic-active enzymes reported so far that show narrow substrate specificity, CsdH showed activity against chemically different polymers with aliphatic (PE and PP) and aromatic (PS) carbon-carbon backbones. Structural analyses, substrate soaking experiments, and biochemical characterisation indicated that the catalytic Cys160-Lys127-Asp39 triad provides a good catalytic environment for accommodating multiple polymer substrates. Moreover, LC-MS identification of degradation products together with SEM, FTIR, and TGA analyses provides supporting evidence for CsdH-mediated true polymer depolymerisation as opposed to simple surface oxidation.

The enzymatic degradation of carbon-carbon backbone polymers is significantly more challenging than that of hydrolysable polyesters such as PET (Suresh et al., 2025). The absence of readily cleavable ester bonds and the highly hydrophobic nature of PE, PP and PS are the main reasons for this (Lewanska and Barczynska, 2025). Thus, previous studies have primarily focused on microbial consortia or whole-cell degradation rather than purified enzymes. For example, marine bacterial communities able to degrade both PET and PE required cooperative interactions between different bacterial species and multiple oxidative enzymes to achieve measurable degradation, highlighting the complexity of C–C polymer biodegradation (Gao and Sun, 2021; Shokunbi et al., 2025; Żarska et al., 2026). Similarly, some bacterial isolates capable of degrading LDPE have exhibited only slight weight loss after extended incubation, implying that polymer oxidation is usually the rate-limiting step in biodegradation (Jebashalomi et al., 2024; Liu et al., 2023). Conversely, the present work identifies a single purified enzyme with broad substrate compatibility, representing an important advance toward enzyme-based recycling strategies.

Recent reviews have pointed out that the identification of enzymes able to degrade polyethylene and polystyrene is still one of the major challenges in plastic biotechnology (Andler et al., 2022; Wei et al., 2025; Żarska et al., 2026). Only a few enzymes have been experimentally characterized, and their catalytic mechanisms are still not well understood (Suresh et al., 2025). The multifunctionality reported for CsdH thus extends the repertoire of plastic-active enzymes known and indicates that cysteine-dependent hydrolases may represent an underexplored class of catalysts for synthetic polymer degradation. The catalytic cysteine residue is more nucleophilic than that of classical serine hydrolases, which may promote oxidation and bond activation on chemically inert polymer surfaces (Cen et al., 2019; Li et al., 1996). The substrate-induced conformational rearrangements observed in the crystal structures provide further evidence that active-site flexibility contributes significantly to substrate promiscuity, allowing accommodation of polymers with different chain architectures.

Degradation products were identified by LC-MS, which further supported the proposed catalytic mechanism. The detection of styrene-derived intermediates and phenylacetic acid during PS degradation suggests oxidative cleavage and metabolism via pathways analogous to bacterial styrene catabolism (Q. Qiu et al., 2024; Yu et al., 2025). Likewise, the detection of hydrocarbon-derived metabolites during PE and PP degradation suggests a stepwise oxidation of long-chain hydrocarbons prior to conversion into lower molecular weight compounds (Sandholm et al., 2025; Ye et al., 2024). These observations are in agreement with the metabolic pathways recently proposed in landfill-derived *Pseudomonas* and *Lysinibacillus* in which the alkane, alcohol dehydrogenases and downstream oxidation enzymes participate in the sequential conversion of oxidised polymer fragments into metabolisable intermediates (Dubey & Thalla, 2025; Jeon et al., 2021; Lv et al., 2024; Wang et al., 2026, 2022).

Our structural analyses provide additional mechanistic insights beyond those previously reported. Crystal structures of the complexes of PE-, PP- and PS-derived substrates and styrene after soaking exhibited the substrate-induced conformational changes in the vicinity of the catalytic pocket. CsdH has an extraordinarily broad substrate spectrum as compared to other characterised cysteine-dependent enzymes. Most cysteine hydrolases are very selective for peptides, amides, thioesters or tiny organic compounds (Lodola et al., 2012; Paulsen and Carroll, 2013), whereas CsdH recognises structurally unrelated molecules of polyethylene, polypropylene, polystyrene and styrene-derived molecules. The catalytic adaptability seen in CsdH consequently enhances the functional diversity of cysteine hydrolases beyond their canonical biological functions. Instead of acting only in peptide or ester hydrolysis (Giles et al., 2003,CsdH seems to be able to start oxidative depolymerisation of very resistant synthetic polymers by combining catalytic nucleophilicity with conformational flexibility. These results highlight cysteine-dependent hydrolase as a novel enzyme family with great promise for building next-generation biocatalysts that can degrade numerous kinds of carbon-carbon backbone polymers simultaneously.

These results further complement recent reports on the novel enzymes involved in polyethylene and polystyrene degradation. Transcriptomic mining of *Stenotrophomonas maltophilia* identified oxidative enzymes such as kynurenine 3-monooxygenase and 4-hydroxyphenylpyruvate dioxygenase as contributors to PS depolymerisation, whereas insect-associated systems have shown more oxidases, hydrolases and dehydrogenases involved in styrene metabolism (Yu et al., 2025). Overall, these studies mainly identified candidate enzymes but did not reveal the structural basis of substrate recognition. The present work addresses this critical knowledge gap by integrating crystallography, biochemical characterisation, and LC-MS validation to describe how a single enzyme can recognise multiple synthetic polymers.

The conservation of the global monomer volume further supports a mechanism of local induced-fit or conformational adaptation, in which the protein architecture remains structurally stable while the substrate-binding cavity dynamically modulates in accordance with the dimensions of the incoming substrate. Taken together, the tunnel and the cavity analyses indicate that substrate promiscuity is achieved through localized structural reorganization rather than global conformational changes. Thus, the enzyme appears to employ a conserved access pathway coupled with a highly adaptable catalytic cavity, providing a structural basis for its ability to accommodate diverse homopolymer hydrocarbon substrates.

Overall, this study represents a major step forward in our understanding of the enzymatic degradation of carbon–carbon backbone homochain plastic polymers. The identification of a multifunctional cysteine hydrolase with activity against PE, PP and PS greatly expands the current repertoire of plastic-active enzymes and demonstrates that structural biology can provide mechanistic insight into broad substrate specificity. These results provide a good basis for rational enzyme engineering for the improvement of catalytic efficiency, substrate affinity and operational stability. In the end, CsdH and its engineered variants could potentially contribute to integrated biological recycling platforms that are able to convert mixed plastic waste into reusable monomers and other value-added products, aiding the development of sustainable circular bioeconomy approaches for plastic waste management

## 5. Conclusion

To summarise, this study presents structural and biochemical evidence that CsdH is a multifunctional cysteine-dependent hydrolase that may operate on the chemically resistant carbon–carbon backbone polymers polyethylene (PE), polypropylene (PP) and polystyrene (PS). Unlike many of the previously characterised plastic-degrading enzymes with tight substrate specificity, CsdH shows broad substrate compatibility for both aliphatic and aromatic polymers. The observed loss of polymer weight, surface morphological changes, FTIR and TGA modifications and identification of low-molecular-weight degradation products by LC–MS imply enzymatic depolymerisation rather than simple surface modification. Further structural investigations showed that the Cys160-Lys127-Asp39 catalytic triad is important for substrate activation and depolymerisation, and the binding-induced conformational changes in the active site aid in the identification of various polymer structures. The crystal structures, and product identification collectively offer a mechanistic basis for the degradation of homopolymers mediated by CsdH. Notably, the discovery of one enzyme interacting with several durable plastics widens the current panel of plastic-active biocatalysts and suggests that cysteine-dependent hydrolases are a largely unexplored class of enzymes for polymer breakdown. These results provide a basis for rational protein engineering to improve catalytic efficiency, substrate affinity and operational stability, and have the long-term potential to develop integrated enzymatic recycling platforms for mixed plastic waste and to promote sustainable circular bioeconomy approaches.

## CRediT authorship contribution statement

**S.A:** Conceptualization, Data analysis, Investigation, Methodology, Writing – original draft, review, and editing; **S.S:** Conceptualization, Data analysis, Investigation, Methodology, Writing – original draft, review, and editing; **A.K:** writing original draft, Data analysis and investigation; **D.A:** Data analysis and investigation, **K.J**: Data analysis and investigation; **M.S**: LCMS analysis, and Writing – original draft, **D.S**: FTIR analysis, **A.K.S:** Investigation, Project administration, Supervision, Writing – review and editing, and **P.K:** Conceptualization, Data curation, Funding acquisition, Investigation, Project administration, Supervision, Writing – review and editing.

## Acknowledgments

The authors thank the Macromolecular Crystallographic Unit (MCU), Institute Instrumentation Centre (IIC), IIT Roorkee, for the X-ray data collection facility. This work was supported by the Ashok Soota Molecular Medicine Facility, Department of Biotechnology, India (DBT), funded projects entitled “Translational and Structural Bioinformatics” – BIC at the Department of Biosciences and Bioengineering, Indian Institute of Technology Roorkee (BT/PR40141/BTIS/137/16/ 2021) and a collaborative project between IIT Roorkee and SKAN Research Trust, Bengaluru. S.A, M.S, D.S, and K.J acknowledge MHRD, A.K. acknowledges DRDO, and D.A. acknowledges UGC, S.S. acknowledges SKAN for financial support.

## Data availability

The complex coordinates of CsdH, along with the structure factors, have been deposited in the RCSB Protein Data Bank (http://www.rcsb.org/pdb) with PDB IDs: 26RX, 44FL, 26RM and 26RB. Other data are available from the corresponding author upon reasonable request.

## Declaration of generative AI

During the preparation of this manuscript, the authors used ChatGPT to improve the language and readability. After using the tool, the authors reviewed and edited the content as needed and take full responsibility for the content of the published article.

## Supplementary Information

**Table S1:**
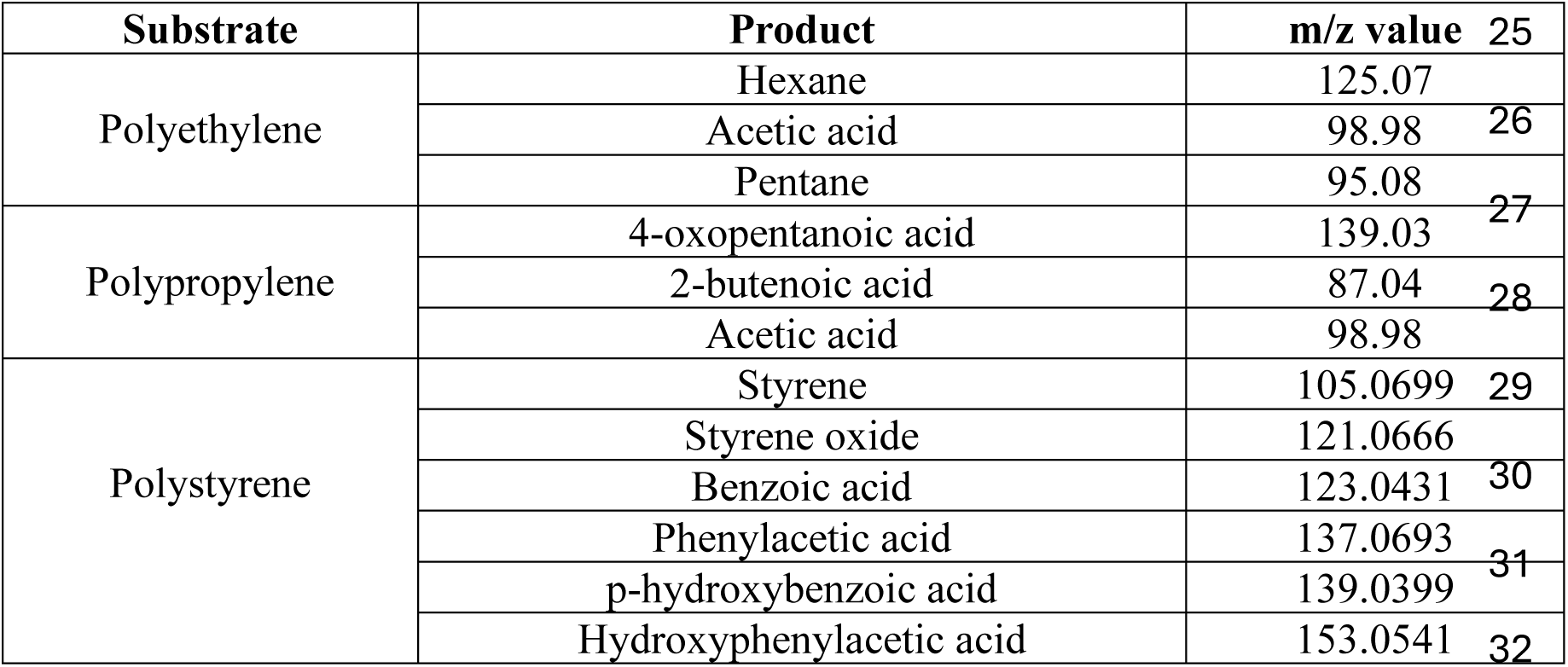
List of substrate and their corresponding degradation products along with their m/z.

**Table S2:** Volumetric comparison of apo and complex structures.

| Structure | Monomer volume (Å<sup>3</sup>) | Active-site cavity (Å<sup>3</sup>) | Increase vs apo |
| --- | --- | --- | --- |
| Apo | 16,482 | 681 | N/A |
| Styrene | 16,719 | 923 | 35.5 % |
| Polystyrene | 16,843 | 1,013 | 48.8 % |
| Polypropylene | 16,308 | 1,392 | 104.4 % |
| Polyethylene | 16,581 | 1,413 | 107.5 % |

**Table S3:** Tunnel analysis of apo-CsdH and complex structures.

| Apo-CsdH |  |  |  |  |  |
| --- | --- | --- | --- | --- | --- |
| Tunnel | Throughput | Cost | Bottleneck radius (Å) | Length (Å) | Curvature |
| 1 | 0.92 | 0.08 | 2.10 | 1.99 | 1.00 |
| 2 | 0.50 | 0.67 | 1.01 | 16.45 | 1.19 |
| CsdH-Styrene complex |  |  |  |  |  |
| Tunnel | Throughput | Cost | Bottleneck radius (Å) | Length(Å) | Curvature |
| 1 | 0.88 | 0.12 | 1.35 | 1.88 | 1.01 |
| 2 | 0.43 | 0.82 | 0.91 | 18.46 | 1.23 |
| CsdH-Polystyrene complex |  |  |  |  |  |
| Tunnel | Throughput | Cost | Bottleneck radius (Å) | Length(Å) | Curvature |
| 1 | 0.89 | 0.11 | 1.80 | 1.46 | 1.06 |
| 2 | 0.51 | 0.66 | 1.08 | 16.95 | 1.20 |
| CsdH-Polypropylene complex |  |  |  |  |  |
| Tunnel | Throughput | Cost | Bottleneck radius (Å) | Length(Å) | Curvature |
| 1 | 0.93 | 0.07 | 2.27 | 1.41 | 1.05 |
| 2 | 0.47 | 0.74 | 0.94 | 14.33 | 1.22 |
| CsdH-Polyethylene complex |  |  |  |  |  |
| Tunnel | Throughput | Cost | Bottleneck radius (Å) | Length(Å) | Curvature |
| 1 | 0.91 | 0.08 | 2.10 | 1.99 | 1.00 |
| 2 | 0.53 | 0.61 | 1.20 | 14.89 | 1.14 |

**Figure S1:**
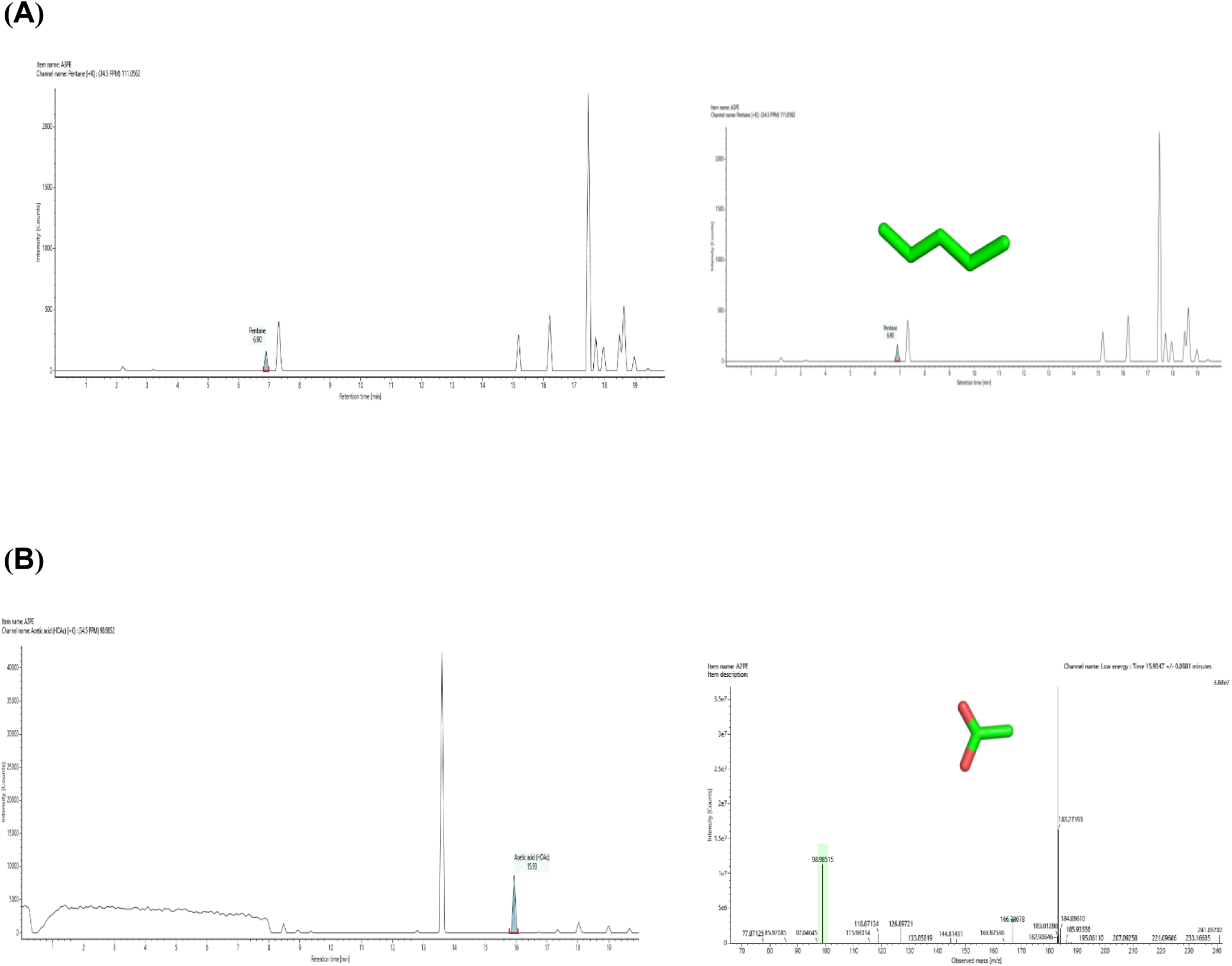
Liquid chromatography–mass spectrometry (LC–MS) analysis of the major PE degradation products generated during enzymatic hydrolysis by CsdH. Representative extracted ion chromatograms (left panels) and corresponding mass spectra (right panels) are shown for **(A)** pentane 6.90 min (m/z 95.08) ([M+ Na]⁺), **(B)** acetic acid 15.93 min (m/z 98.98) ([M+ K]⁺).

**Figure S2:**
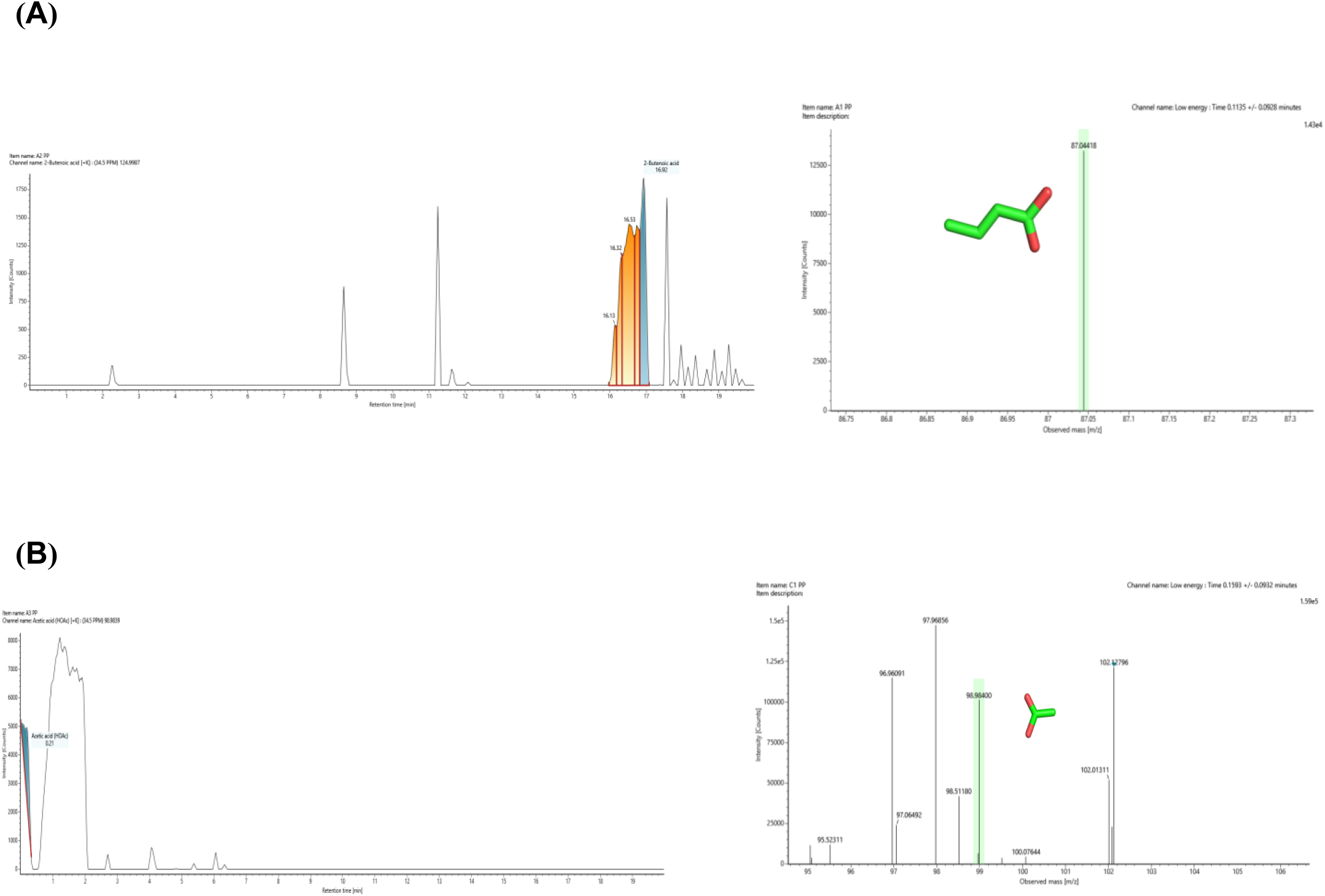
Liquid chromatography–mass spectrometry (LC–MS) analysis of the major PP degradation products generated during enzymatic hydrolysis by CsdH. Representative extracted ion chromatograms (left panels) and corresponding mass spectra (right panels) are shown for **(A)** 2-butenoic acid 17.07 min (m/z 87.04) ([M+ H]⁺), and **(B)** acetic acid 15.93 min (m/z 98.98) ([M+ K]⁺).

**Figure S3:**
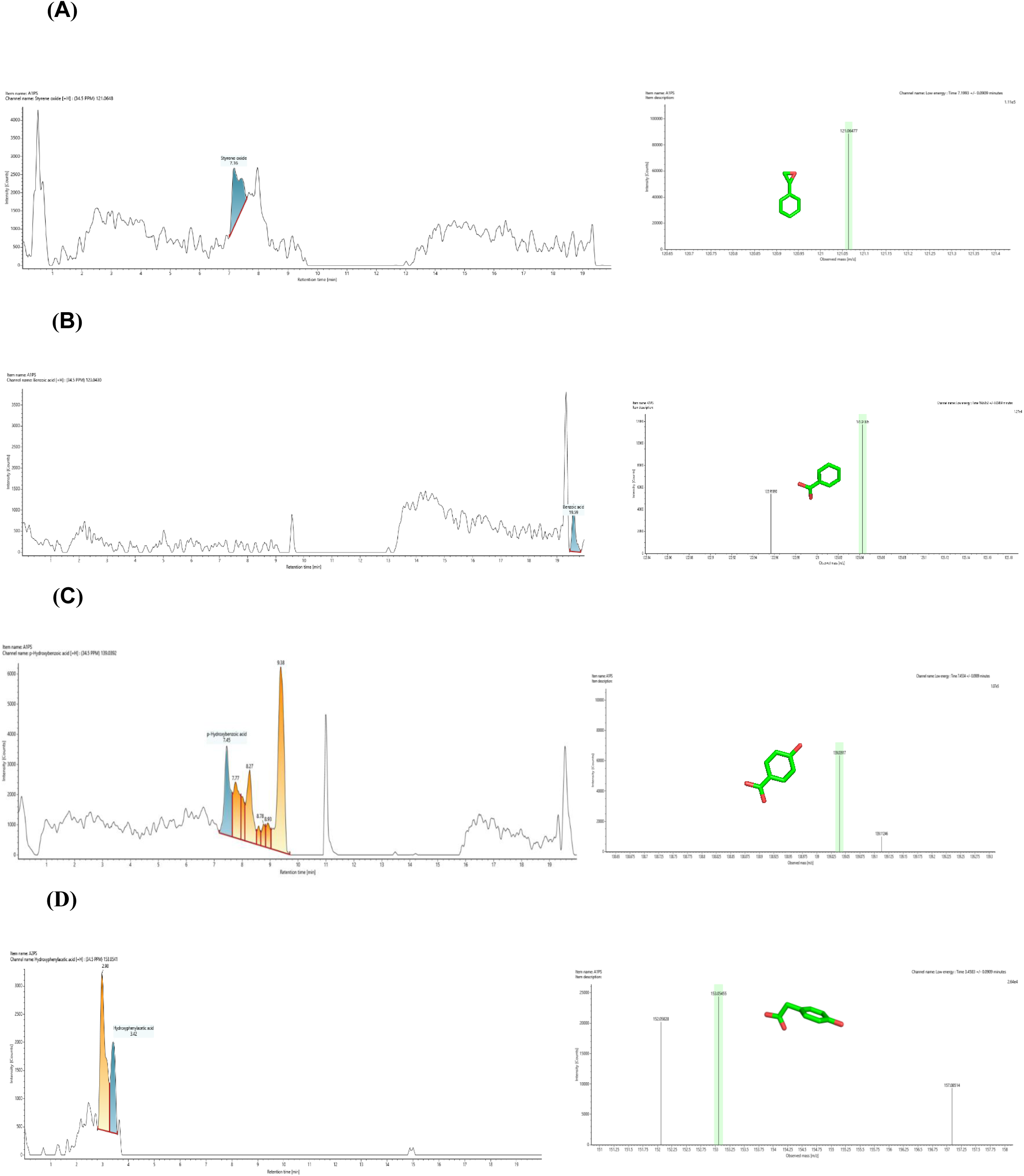
Liquid chromatography–mass spectrometry (LC–MS) analysis of the major PE, PP, and degradation products generated during enzymatic hydrolysis by CsdH. Representative extracted ion chromatograms (left panels) and corresponding mass spectra (right panels) are shown for **(A)** styrene oxide 7.16 min (m/z 121.0666) ([M+ H]⁺), **(B)** benzoic acid 19.51 min (m/z 123.0431) ([M+ H]⁺), **(C)** p-hydroxybenzoic acid 7.45 min (m/z 137.0693) ([M+ H]⁺), and **(D)** p-hydroxyphenylacetic acid 3.42 min (m/z 153.0541) ([M+ H]⁺)

**Figure S4:**
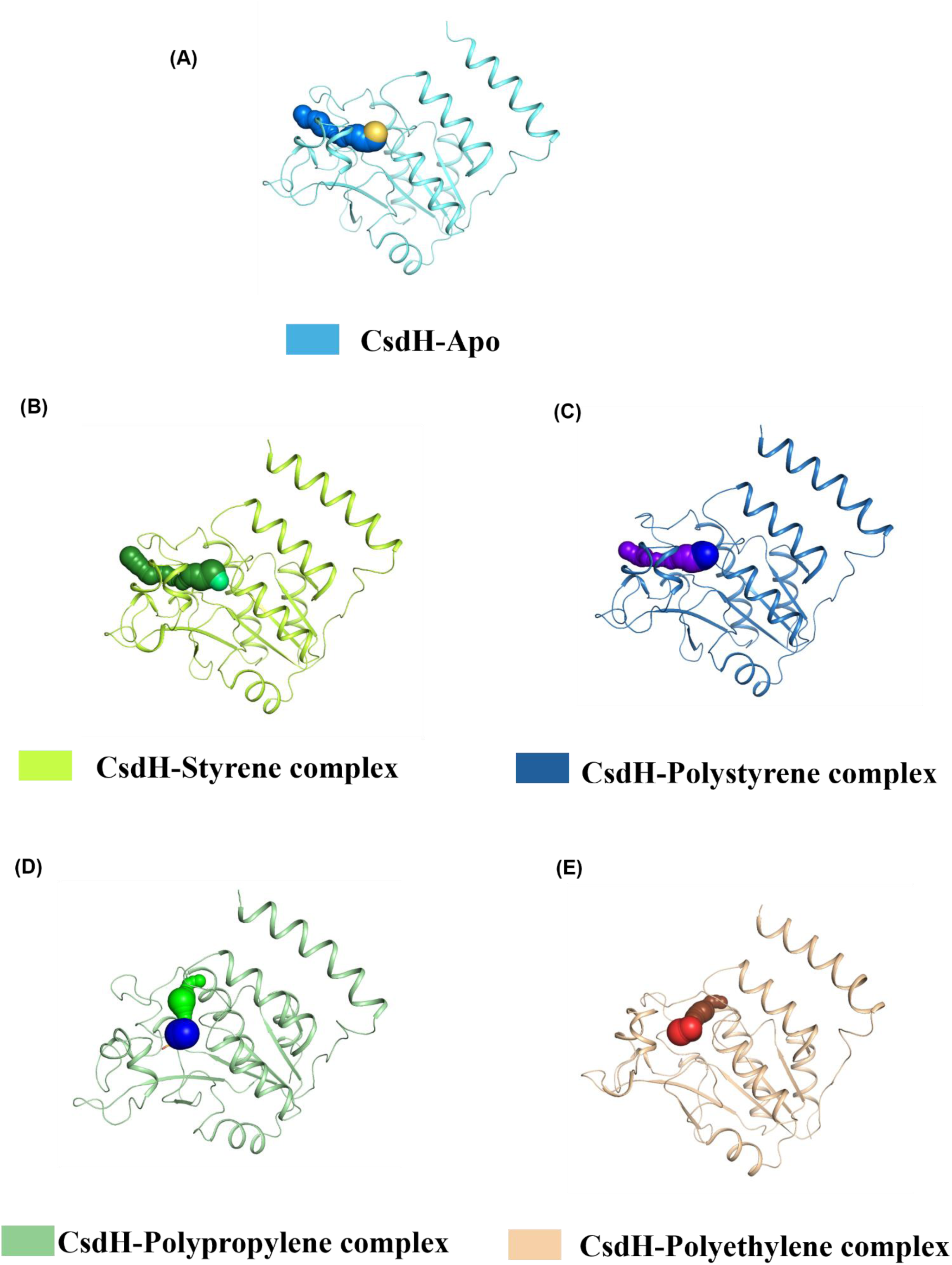
Tunnel and cavity analysis of **(A)** apo-CsdH **(B)** styrene complex **(C)** polystyrene complex **(D)** polypropylene complex **(E)** polyethylene complex

## References

Aggarwal, S., Singh, S., Jangid, K., Aggarwal, D., Kayastha, A., Shakib, M., Sharma, A.K., Kumar, P., 2026. Discovery and Structural Insights into a Cysteine-Dependent PET Hydrolase. 10.2139/SSRN.7247458

Alharbi, M., Abd-elhaliem, S.M., Afifi, S.S., Al-shareef, walaa A., Mosbah, R.A., Mansour, A.T., Alharbi, N.K., Bendary, M.M., 2026. Toward sustainable plastic bioremediation using bacterial consortia from aquatic environments. Front. Bioeng. Biotechnol. 13, 1709072. 10.3389/FBIOE.2025.1709072/TEXT

Andler, R., Tiso, T., Blank, L., Andreeßen, C., Zampolli, J., D’Afonseca, V., Guajardo, C., Díaz-Barrera, A., 2022. Current progress on the biodegradation of synthetic plastics: from fundamentals to biotechnological applications. Reviews in Environmental Science and Bio/Technology 2022 21:4 21, 829–850. 10.1007/S11157-022-09631-2

Appel, M.J., Bertozzi, C.R., 2015. Formylglycine, a Post-Translationally Generated Residue with Unique Catalytic Capabilities and Biotechnology Applications. ACS Chem. Biol. 10, 72. 10.1021/CB500897W

Bher, A., Mayekar, P.C., Auras, R.A., Schvezov, C.E., 2022. Biodegradation of Biodegradable Polymers in Mesophilic Aerobic Environments. International Journal of Molecular Sciences 2022, Vol. 23, Page 12165 23, 12165. 10.3390/IJMS232012165

Cen, Y., Singh, W., Arkin, M., Moody, T.S., Huang, M., Zhou, J., Wu, Q., Reetz, M.T., 2019. Artificial cysteine-lipases with high activity and altered catalytic mechanism created by laboratory evolution. Nat. Commun. 10. 10.1038/s41467-019-11155-3

Chatonnet, A., Perochon, M., Velluet, E., Marchot, P., 2023. The ESTHER database on alpha/beta hydrolase fold proteins - An overview of recent developments. Chem. Biol. Interact. 383, 110671. 10.1016/J.CBI.2023.110671

Chovancova, E., Pavelka, A., Benes, P., Strnad, O., Brezovsky, J., Kozlikova, B., Gora, A., Sustr, V., Klvana, M., Medek, P., Biedermannova, L., Sochor, J., Damborsky, J., 2012. CAVER 3.0: A Tool for the Analysis of Transport Pathways in Dynamic Protein Structures. PLoS Comput. Biol. 8, e1002708. 10.1371/JOURNAL.PCBI.1002708

Danso, D., Chow, J., Streita, W.R., 2019. Plastics: Environmental and biotechnological perspectives on microbial degradation. Appl. Environ. Microbiol. 85. 10.1128/AEM.01095-19;CTYPE:STRING:JOURNAL

Delano, W.L., 2002. Delano, W.L. (2002) An Open-Source Molecular Graphics Tool. https://www.scirp.org/reference/ReferencesPapers?ReferenceID=2323923 (accessed 8.18.26).

Dubey, A.P., Thalla, A.K., 2025. Bioprospecting indigenous bacteria from landfill leachate for enhanced polypropylene microplastics degradation. J. Hazard. Mater. 487, 137139. 10.1016/J.JHAZMAT.2025.137139

Gao, R., Sun, C., 2021. A marine bacterial community capable of degrading poly(ethylene terephthalate) and polyethylene. J. Hazard. Mater. 416, 125928. 10.1016/J.JHAZMAT.2021.125928

Giles, N.M., Giles, G.I., Jacob, C., 2003. Multiple roles of cysteine in biocatalysis. Biochem. Biophys. Res. Commun. 300, 1–4. 10.1016/S0006-291X(02)02770-5

Hamer, G., 2003. Solid waste treatment and disposal: effects on public health and environmental safety. Biotechnol. Adv. 22, 71–79. 10.1016/J.BIOTECHADV.2003.08.007

Ho, B.T., Roberts, T.K., Lucas, S., 2018. An overview on biodegradation of polystyrene and modified polystyrene: the microbial approach. Crit. Rev. Biotechnol. 38, 308–320. 10.1080/07388551.2017.1355293

Hossain, M., Engelhardt, I., 2025. Global plastic footprint: unveiling property trends, environmental fate, and emerging threats of microplastic and nanoplastics pollution across ecosystems. Energy, Ecology and Environment 2025 10:6 10, 637–674. 10.1007/S40974-025-00383-7

Jebashalomi, V., Emmanuel Charles, P., Rajaram, R., 2024. Microbial degradation of low-density polyethylene (LDPE) and polystyrene using Bacillus cereus (OR268710) isolated from plastic-polluted tropical coastal environment. Science of The Total Environment 924, 171580. 10.1016/J.SCITOTENV.2024.171580

Jeon, J.M., Park, S.J., Choi, T.R., Park, J.H., Yang, Y.H., Yoon, J.J., 2021. Biodegradation of polyethylene and polypropylene by Lysinibacillus species JJY0216 isolated from soil grove. Polym. Degrad. Stab. 191, 109662. 10.1016/J.POLYMDEGRADSTAB.2021.109662

Kayastha, A., Jangid, K., Verma, S., Rana, A., Kumar, P., 2026. Structural and mechanistic insights into Edo4, an extradiol dioxygenase in dioxin degradation. Arch. Biochem. Biophys. 782, 110878. 10.1016/J.ABB.2026.110878

Kunwar, B., Cheng, H.N., Chandrashekaran, S.R., Sharma, B.K., 2016. Plastics to fuel: a review. Renewable and Sustainable Energy Reviews 54, 421–428. 10.1016/J.RSER.2015.10.015

Lewanska, M., Barczynska, R., 2025. Microplastics from Food Packaging: Polymer Degradation Pathways, Environmental Distribution, and Effects on the Human Gastrointestinal Tract. Polymers 2025, Vol. 17, Page 2923 17, 2923. 10.3390/POLYM17212923

Li, J., Szittner, R., Derewenda, Z.S., Meighen, E.A., 1996. Articles Conversion of Serine-114 to Cysteine-114 and the Role of the Active Site Nucleophile in Acyl Transfer by Myristoyl-ACP Thioesterase from Vibrio harVeyi †.

Liu, R., Zhao, S., Zhang, B., Li, G., Fu, X., Yan, P., Shao, Z., 2023. Biodegradation of polystyrene (PS) by marine bacteria in mangrove ecosystem. J. Hazard. Mater. 442, 130056. 10.1016/J.JHAZMAT.2022.130056

Liu, W., Lu, L., Zeng, X., Zhang, T., Ma, L., 2026. Microbial synergism in a synthetic consortium enabled efficient polyethylene biodegradation. Ecotoxicol. Environ. Saf. 316, 120151. 10.1016/J.ECOENV.2026.120151

Lodola, A., Branduardi, D., de Vivo, M., Capoferri, L., Mor, M., Piomelli, D., Cavalli, A., 2012. A catalytic mechanism for cysteine N-terminal nucleophile hydrolases, as revealed by free energy simulations. PLoS One 7. 10.1371/journal.pone.0032397

Lv, S., Li, Y., Zhao, S., Shao, Z., 2024. Biodegradation of Typical Plastics: From Microbial Diversity to Metabolic Mechanisms. International Journal of Molecular Sciences 2024, Vol. 25, Page 593 25, 593. 10.3390/IJMS25010593

Macheca, A.D., Mutuma, B., Adalima, J.L., Midheme, E., Lúcas, L.H.M., Ochanda, V.K., Mhlanga, S.D., 2024. Perspectives on Plastic Waste Management: Challenges and Possible Solutions to Ensure Its Sustainable Use. Recycling 2024, Vol. 9, Page 77 9, 77. 10.3390/RECYCLING9050077

Mahto, J.K., Mishra, I., Jangid, K., Kumar, P., 2025. Structure-guided engineering of an aromatic ring-hydroxylating dioxygenase for broad-spectrum phthalate degradation. J. Bacteriol. 207. 10.1128/JB.00221-25;PAGE:STRING:ARTICLE/CHAPTER

Mahto, J.K., Sharma, M., Neetu, N., Kayastha, A., Aggarwal, S., Kumar, P., 2022. Conformational flexibility enables catalysis of phthalate cis-4,5-dihydrodiol dehydrogenase. Arch. Biochem. Biophys. 727. 10.1016/J.ABB.2022.109314

Maity, W., Maity, S., Bera, S., Roy, A., 2021. Emerging Roles of PETase and MHETase in the Biodegradation of Plastic Wastes. Applied Biochemistry and Biotechnology 2021 193:8 193, 2699–2716. 10.1007/S12010-021-03562-4

Mamtimin, T., Han, H., Khan, A., Feng, P., Zhang, Q., Ma, X., Fang, Y., Liu, P., Kulshrestha, S., Shigaki, T., Li, X., 2023. Gut microbiome of mealworms (Tenebrio molitor Larvae) show similar responses to polystyrene and corn straw diets. Microbiome 11, 1–20. 10.1186/S40168-023-01550-W/FIGURES/3

Meran, C., Ozturk, O., Yuksel, M., 2008. Examination of the possibility of recycling and utilizing recycled polyethylene and polypropylene. Mater. Des. 29, 701–705. 10.1016/J.MATDES.2007.02.007

Nikolaivits, E., Pantelic, B., Azeem, M., Taxeidis, G., Babu, R., Topakas, E., Brennan Fournet, M., Nikodinovic-Runic, J., 2021. Progressing Plastics Circularity: A Review of Mechano-Biocatalytic Approaches for Waste Plastic (Re)valorization. Front. Bioeng. Biotechnol. 9, 696040. 10.3389/FBIOE.2021.696040/FULL

Oanca, G., Asadi, M., Saha, A., Ramachandran, B., Warshel, A., 2020. Exploring the Catalytic Reaction of Cysteine Proteases. J. Phys. Chem. B 124, 11349–11356. 10.1021/ACS.JPCB.0C08192

Paulsen, C.E., Carroll, K.S., 2013. Cysteine-Mediated Redox Signaling: Chemistry, Biology, and Tools for Discovery. Chem. Rev. 113, 4633. 10.1021/CR300163E

Pérez-García, P., Sass, K., Wongwattanarat, S., Amann, J., Feuerriegel, G., Neumann, T., Bäse, N., Schmitz, L.S., Dierkes, R.F., Gurschke, M.F., Wypych, A., Bounabi, H., de Divitiis, M., Vollstedt, C., Streit, W.R., 2025. Microbial plastic degradation: enzymes, pathways, challenges, and perspectives. Microbiology and Molecular Biology Reviews 89. 10.1128/MMBR.00087-24;JOURNAL:JOURNAL:BR;PAGE:STRING:ARTICLE/CHAPTER

Qiu, J., Chen, Y., Zhang, L., Wu, J., Zeng, X., Shi, X., Liu, L., Chen, J., 2024. A comprehensive review on enzymatic biodegradation of polyethylene terephthalate. Environ. Res. 240, 117427. 10.1016/J.ENVRES.2023.117427

Qiu, Q., Li, H., Sun, X., Tian, K., Gu, J., Zhang, F., Zhou, D., Zhang, X., Huo, H., 2024. Integrating genomics, molecular docking, and protein expression to explore new perspectives on polystyrene biodegradation. J. Hazard. Mater. 476, 135031. 10.1016/J.JHAZMAT.2024.135031

Raček, T., Vel’Ký, D., Bučeková, G., Schindler, O., Hutařová Vařeková, I., Špačková, A., Bazgier, V., Berka, K., Svobodová, R., 2025. MOLEonline: a web-based tool for analysing channels, tunnels, and pores (2025 update). Bioinformatics 41. 10.1093/BIOINFORMATICS/BTAF486

Sandholm, R.M., Boehlich, G.J., Dahl, Ø., Chowreddy, R.R., Stepnov, A., Vaaje-Kolstad, G., La Rosa, S.L., 2025. Microbial degradation of a widely used model polyethylene is restricted to medium- and long-chain alkanes and their oxidized derivatives. ISME J. 19. 10.1093/ISMEJO/WRAF276

Shehbas, M.C., Desai, D.S., Mathew, T.M., Nampoothiri, K.M., 2026. Microbial enzymes for plastic degradation: a comprehensive review of current status and emerging trends. Biodegradation 2026 37:2 37, 30-. 10.1007/S10532-026-10248-3

Shokunbi, A.O., Ezeh, O.M., Fadipe, D.P., Olanrewaju, A.S., Badru, O.O., Eze, C.N., Iwuala, L.C., 2025. Microbial Consortia: Synergistic Effects on Plastic Degradation and Enzyme Production. Journal of Medical Science, Biology, and Chemistry 2, 149–156. 10.69739/JMSBC.V2I1.681

Singh, S., Bajaj, A., Manickam, N., 2026. Microbiome of soil waste dumpsite and adjacent river habitat harbors dynamic plastic degrading bacterial diversity and abundant functional enzymes. Science of The Total Environment 1014, 181331. 10.1016/J.SCITOTENV.2025.181331

Singh, S., Soni, M., Gupta, N., Sandhu, P., Tripathi, D., Venkatesh Pratap, J., Subramanian, S., Manickam, N., 2024a. Unravelling biochemical and molecular mechanism of a carboxylesterase from Dietzia kunjamensis IITR165 reveal novel activities against polyethylene terephthalate. Biochem. Biophys. Res. Commun. 735, 150833. 10.1016/J.BBRC.2024.150833

Singh, S., Subramanian, S., Gupta, N., Bajaj, A., Manickam, N., 2024b. Genomic insights on gene clusters and pathways for the biodegradation of plastic compounds: Unravelling the metabolic versatility in a Dietzia kunjamensis IITR165. Curr. Res. Biotechnol. 8, 100258. 10.1016/J.CRBIOT.2024.100258

Singh, S., Thakur, R.S., Manickam, N., 2023. Insights into molecular mechanism of plasticizer biodegradation in Dietzia kunjamensis IITR165 and Brucella intermedia IITR166 isolated from a solid waste dumpsite. J. Appl. Microbiol. 134. 10.1093/JAMBIO/LXAD231

Son, H.F., Cho, I.J., Joo, S., Seo, H., Sagong, H.Y., Choi, S.Y., Lee, S.Y., Kim, K.J., 2019. Rational Protein Engineering of Thermo-Stable PETase from Ideonella sakaiensis for Highly Efficient PET Degradation. ACS Catal. 9, 3519–3526. 10.1021/ACSCATAL.9B00568

Srivastava, P., Saji, J., Manickam, N., 2024. Biodegradation of polyethylene terephthalate (PET) by Brucella intermedia IITR130 and its proposed metabolic pathway. Biodegradation 2024 35:5 35, 671–685. 10.1007/S10532-024-10070-9

Srivastava, P., Singh, S., Soni, M., Pratap, J.V., Subramanian, S., Manickam, N., 2025. Enzymatic degradation of PET by hydrolase from Brucella intermedia IITR130 and its genomic insights. Biodegradation 36, 1–20. 10.1007/S10532-025-10141-5/METRICS

Suresh, V., Shams, R., Dash, K.K., Shaikh, A.M., Béla, K., 2025. Comprehensive review on enzymatic polymer degradation: A sustainable solution for plastics. J. Agric. Food Res. 20, 101788. 10.1016/J.JAFR.2025.101788

Thadepalli, S., 2022. Review of multifarious applications of polymers in medical and health care textiles. Mater. Today Proc. 55, 330–336. 10.1016/J.MATPR.2021.07.513

Verma, S., Aggarwal, D., Ashar, M., Pandey, A.K., Sutradhar, A., Pandey, S., Sircar, D., Singla, J., Kumar, P., 2026. Structural and functional characterization of thermostable EstS1 esterase for BHET degradation. J. Struct. Biol. 218, 108342. 10.1016/J.JSB.2026.108342

Wang, D., Yang, Y., Ye, Y., He, D., Fu, Q., Liu, X., 2026. Microbial Degradation of Plastics: Mechanisms, Pathways, and Multiomics Insights. Environ. Sci. Technol. 60, 17076–17098. 10.1021/ACS.EST.6C02515

Wang, P., Zhao, J., Ruan, Y., Cai, X., Li, J., Zhang, L., Huang, H., 2022. Degradation of Polypropylene by the Pseudomonas aeruginosa Strains LICME WZH-4 and WGH-6. Journal of Polymers and the Environment 2022 30:9 30, 3949–3958. 10.1007/S10924-022-02480-8

Wei, R., Weber, G., Blank, L.M., Bornscheuer, U.T., 2025. Process insights for harnessing biotechnology for plastic depolymerization. Nature Chemical Engineering 2025 2:2 2, 110–117. 10.1038/s44286-024-00171-w

Ye, J., Liu, H., Gao, W., Li, N., Cheng, Z., Yan, B., Chen, G., 2024. Radicals mediated polypropylene waste plastics oxidation into ketone fuel. Chemical Engineering Journal 490, 151899. 10.1016/J.CEJ.2024.151899

Yu, Y., Tian, K., Hao, P., Wang, Y., Chang, M., Zhou, D., Gu, J., Zhang, F., Huo, H., 2025. Enzymatic mechanism of polystyrene biodegradation: key depolymerization enzymes in the Stenotrophomonas maltophilia sp. ZSL2493. Environ. Res. 285, 122659. 10.1016/J.ENVRES.2025.122659

Żarska, M.J., Jasiak, M.D., Mierzejewski, P., Tomczyk, K., Małecki, J., Gudz, R., Urbanek, A.K., Kosiorowska, K.E., Dybka, J.A., 2026. Plastic-Degrading Microorganisms: Biodegradation Pathways and Habitat Origins. Molecules 31, 1638. 10.3390/MOLECULES31101638/S1

